# Serotype switching in *Pseudomonas aeruginosa* ST111 enhances adhesion and virulence

**DOI:** 10.1101/2024.04.26.591252

**Authors:** Mikkel Anbo, Mahbuba Akter Lubna, Dina A. Moustafa, Telmo O. Paiva, Laura Serioli, Kinga Zor, Claus Sternberg, Katy Jeannot, Yves F. Dufrêne, Joanna B. Goldberg, Lars Jelsbak

## Abstract

Evolution of the highly successful and multidrug resistant clone ST111 in *Pseudomonas aeruginosa* involves serotype switching from O-antigen O4 to O12. How expression of a different O-antigen serotype alters pathogen physiology to enable global dissemination of this high-risk clone-type is not understood. Here, we engineered isogenic laboratory and clinical *P. aeruginosa* strains that express the different O-antigen gene clusters to assess the correlation of structural differences of O4 and O12 O-antigens to pathogen-relevant phenotypic traits. We show that serotype O12 is associated with enhanced adhesion, type IV pili dependent twitching motility, and tolerance to host defense molecules and serum. Moreover, we find that serotype O4 is less virulent compared to O12 in an acute murine pneumonia infection in terms of both colonization and survival rate. Finally, we find that these O-antigen effects may be explained by specific biophysical properties of the serotype repeat unit found in O4 and O12, and by differences in membrane stability between O4 and O12 expressing cells. The results demonstrate that differences in O-antigen sugar composition can directly affect *P. aeruginosa* pathogenicity traits, and provide a better understanding of the potential selective advantages that underlie serotype switching and emergence of serotype O12 ST111.

## Introduction

The increasing prevalence of nosocomial infections caused by multidrug-resistant (MDR) or extensively drug-resistant (XDR) *Pseudomonas aeruginosa* is frequently linked to a limited number of so-called epidemic ‘high-risk cloneś (HiRiCs) that are widespread in hospitals worldwide (1, 2). HiRiCs are associated with poor clinical outcomes (1, 2) and contain a large and diverse number of horizontally acquired resistance determinants (3), yet the molecular mechanisms that enable particular clone types (but not others) to emerge as HiRiCs remain poorly understood.

We have previously shown that evolution of one of the most globally disseminated HiRiCs, ST111 serotype O12, is associated with a serotype switch in which the native O-antigen O4 biosynthesis gene cluster was replaced by the O12 gene cluster by horizontal gene transfer and recombination (4). Although ST111 is globally associated both with serotype O12 and O4, ST111 O12 is by far more prevalent than ST111 O4 among antibiotic resistant clinical isolates (5–8), suggesting that the switch from O4 to O12 expression provides ST111 with unknown fitness advantages. In support of this, we have also observed multiple, independent serotype switches from O4 to O12 in ST111 isolates across different continents (4, 9), and thus it appears that there is a strong selection for the recombinatorial replacement of O4 with O12 in ST111. Other observations also point towards an important role of serotype O12 in additional clone types. For example, we have previously observed - albeit sporadically – serotype switches from other serotypes to O12 in other clone types, such as ST253 (PA14-like, usually associated with serotype O19) and ST244 (usually associated with serotype O2/O5) (9). Furthermore, serotype O12 strains have been associated with multidrug resistance in Europe since the 1980s (10). The aim of the present study was to determine how the switch from serotype O4 to O12 alters pathogen behavior, fitness or virulence of ST111 to explain the selective advantage of the serotype switch and the global success of this devastating clone type.

The lipopolysaccharide (LPS) is a major virulence factor of *P. aeruginosa* which has been shown to interact with myriads of molecular species, including host molecules, antibiotics, and bacteriophages (11–16). LPS is composed of the lipid A, a core polysaccharide, and a variable O-specific antigen (OSA) or the common polysaccharide antigen (CPA). The O-antigen is composed of repeat units of 2-5 sugars and their chemical structures, and the underlying genetics have been described previously (12, 17). The variable and diverse structures of repeating sugar subunits in OSA classify *P. aeruginosa* into 20 distinct serotypes (18–20). By the disruption of O-antigen synthesis through genetic knockouts, many studies have previously documented the importance of the O-antigen to clinically relevant phenotypes such as virulence in models of acute infection (21), antibiotic susceptibility (15, 16, 22), and host complement-mediated killing (23). Although these findings clearly illustrate the general importance of the O-antigen structure, we currently do not understand how the different sugar compositions found in the various O-antigens expressed on *P. aeruginosa* affect the biology of the pathogen. The serotype switch documented in ST111 strongly suggests that different O-antigen structures may indeed have different, yet unknown, effects on *P. aeruginosa* biology and virulence. Similarly, studies have shown that a number of serotypes predominate in clinical settings and epidemic outbreaks (24), suggesting a link between serotypes and pathogen behavior and physiology. However, these epidemiological studies do not offer straightforward explanations as to why particular serotypes would predominate and how different serotypes affect pathogen biology.

Associating particular phenotypes to specific O-antigen structures by comparing clinical isolates of different serotypes is not straightforward and confounded by the genetic diversity of the different *P. aeruginosa* hosts. To overcome this problem, we constructed isogenic strains that express either O4 or O12 serotype with the goal of defining how these different O-antigen structures affect *P. aeruginosa* physiology. Here, we show that adhesion properties and virulence phenotypes are differentially affected by heterologous expression of O4 and O12 in different *P. aeruginosa* backgrounds, specifically we noted that O12 is more adherent and virulent in our models. We furthermore identify serotype-specific alterations in membrane stability and function of membrane associated proteins as the mechanism underlying the adhesion and virulence phenotypes. These results may help to explain the global clonal success of multi-drug resistant ST111 serotype O12.

## Materials and methods

### Bacterial strains, plasmids and culture conditions

Bacterial strains and plasmids used in this study are described in Table 1 and 2, respectively. *P. aeruginosa* and *E. coli* overnight cultures were routinely grown in LB for 16-18 hours at 37°C, 200 rpm unless stated otherwise. Cells were stored at -80°C in a 20% glycerol solution and streaked on LB agar (2%) for further sub-culturing. For selection or maintenance of plasmids in *E. coli* the following supplements were added: 8 µg/ml tetracycline, 100 µg/ml ampicillin, 35 µg/ml kanamycin, 10 µg/ml gentamycin, 6 µg/ml chloramphenicol. For selection of *P. aeruginosa* the following supplements were added: 50-100 µg/ml tetracycline, 50 µg/ml gentamycin. Competent *E. coli* cells were prepared by repeat washing with 100mM CaCl_2_ + 15% glycerol and 50 µl aliquots were transformed by addition of plasmid and heat shock (30s at 42°C, 2 min on ice, rescued for 1-2 h at 37°C after addition of 1 ml LB). Expression of O-antigen was verified by agglutination with *Pseudomonas aeruginosa* monovalent anti-serum P4, P5, P10 and P12 (Bio-Rad).

**Table 1.** Bacterial strains used in this study. *Abbreviations used: ETγA for genes recE, recT, redγ, and recA. OSA for O-specific antigen.

| Name | Description* | Source |
| --- | --- | --- |
| ***E. coli* strains** |  |  |
| Cc118λpir | Plasmid propagation | (25) |
| GBdir-pir116 | Arabinose inducible ETyA operon | (26) |
| ***P. aeruginosa* strains** |  |  |
| PAO1 | Laboratory strain | ATCC15692 |
| PA14 | Laboratory strain | Janus Haagesen, DTU Biosustain |
| ST111 2875 | Strain 2875, serotype O4 | (27) |
| ST111 2879 | Strain 2879, serotype O12 | (27) |
| IATS O4 | Serotype O4 type strain | Joseph Lam |
| IATS O12 | Serotype O12 type strain | Joseph Lam |
| PAO1ΔO | PAO1 Δ*wbpM*1741-3547432 (deletes OSA) | This study |
| PA14ΔO | PA14 Δ2026751-*wbpM*1741 (deletes OSA) | This study |
| ST111ΔO | ST111 2875 Δ2113766-*wbpM*1741 (deletes OSA) | This study |
| PA14ΔO+O4 | PA14 ΔO attTn7::O4 | This study |
| PA14ΔO+O12 | PA14 ΔO attTn7::O12 | This study |
| PA14ΔO+O19 | PA14 ΔO attTn7::O19 | This study |
| PAO1ΔO+O4 | PAO1 ΔO attTn7::O4 | This study |
| PAO1ΔO+O12 | PAO1 ΔO attTn7::O12 | This study |
| PAO1ΔO+O5 | PAO1 ΔO attTn7::O5 | This study |
| ST111ΔO+O4 | ST111 ΔO attTn7::O4 | This study |
| ST111ΔO+O12 | ST111 ΔO attTn7::O12 | This study |
| PAO1 galU | Rough LPS mutant | (28) |
| PAO1pilA | Type IV pili deficient mutant | (29) |

**Table 2.** Plasmids used in this study. *Abbreviations and nomenclature: OSA for O-specific antigen. eYFP for enhanced yellow fluorescent protein. msfGFP for monomeric superfolder green fluorescent protein. mKate is a red fluorescent protein. CamR for chloramphenicol resistant. AmpR for ampicillin resistant. SacB is the Bacillus subtilis levansucrase gene. TetR for tetracycline resistant. KmR for kanamycin resistant, and GmR for gentamycin resistant.

| Name | Description* | Selection | Source |
| --- | --- | --- | --- |
| pRK600 | Conjugation helper plasmid | CamR | (30) |
| pTNS2 | Tn7 helper plasmid | AmpR | (31) |
| pFLP3 | Flp expression vector | SacB AmpR<br>TetR | (31) |
| pNJ1 | Allelic replacement | TetR SacB | (32) |
| pMA03 PAO1 | pNJ1 derivate,<br>facilitates deletion of<br>OSA cluster in PAO1 | TetR SacB | This study |
| pMA03 PA14 | pNJ1 derivate,<br>facilitates deletion of<br>OSA cluster in PA14 | TetR SacB | This study |
| pMA03 ST111 | pNJ1 derivate,<br>facilitates deletion of<br>OSA cluster in ST111 | TetR SacB | This study |
| pMA10 PAO1 | pNJ1 derivate,<br>facilitates integration<br>of *tetA* in serotype<br>cluster of PAO1 | TetR SacB | This study |
| pMA10 PA14 | pNJ1 derivate,<br>facilitates integration<br>of *tetA* in serotype<br>cluster of PA14 | TetR SacB | This study |
| pMA10 O4 | pNJ1 derivate,<br>facilitates integration<br>of *tetA* in serotype<br>cluster of IATS O4 | TetR SacB | This study |
| pMA10 O12 | pNJ1 derivate,<br>facilitates integration<br>of *tetA* in serotype<br>cluster of IATS O12 | TetR SacB | This study |
| pUC18R6kT-mini-Tn7T-Km | Tn7 tagging vector with R6K oriR incP oriT | AmpR, KmR | (31) |
| Mini-tn7(GM) PA<sub>A1/04/03</sub>-eyfp-A | Tn7 tagging vector with eYFP | GmR, CamR, AmpR, eYFP | (33) |
| pMA09 | Mini-tn7 derivate, with eYFP integrated between two transcriptional terminators | AmpR, KmR, eYFP | This study |
| pMA11 O5 | pMA09 derivate, with PAO1 serotype cluster (O5) replacing eYFP | TetR, KmR, AmpR, SacB | This study |
| pMA11 O19 | pMA09 derivate, with PA14 serotype cluster (O19) replacing eYFP | TetR, KmR, AmpR, SacB | This study |
| pMA11 O4 | pMA09 derivate, with IATS O4 serotype cluster (O4) replacing eYFP | TetR, KmR, AmpR, SacB | This study |
| pMA11 O12 | pMA09 derivate, with IATS O12 serotype cluster replacing eYFP | TetR, KmR, AmpR, SacB | This study |
| pBG42 msfGFP | Tn7 construct expressing msfGFP | GmR | (34) |
| pBG42 mKate | Tn7 construct expressing mKate | GmR | Hansen, M unpublished |
| Mini-ctx2 | Genomic tagging vector | TetR | (35) |
| Mini-ctx2 msfGFP<br>aacC1 | Mini-ctx2 derivate<br>expressing msfGFP<br>and *aacC1* | TetR GmR | This study |
| Mini-ctx2 mKate<br>aacC1 | Mini-ctx2 derivate<br>expressing mKate and<br>*aacC1* | TetR GmR | This study |
| pSEVA658::pilA<sub>PA14</sub> | pSEVA658<br>overexpression vector<br>for pilA (from PA14),<br>inducible by addition<br>of m-toluate | Gm | This study |

### Bacterial growth media

For bacterial growth, the following culture media were used: Lysogeny broth (LB; Carl Roth, X964), Pseudomonas isolation agar (PIA; Millipore, 17208), Low salt LB (LSLB; 10 g/l tryptone, 5 g/l yeast extract, 0.5 g/l NaCl) and no salt LB agar (NSLB; 10 g/l tryptone, 5 g/l yeast extract, 15 g/l agar, 6% sucrose). For solid growth media, agar (ITW Reagents, A0949) was supplemented at 2% (w/v) unless otherwise stated. Cultures were regularly washed with PBS (8 g/l NaCl, 0.2 g/l KCl, 1.44 g/l Na_2_HPO_4_·2H_2_O, 0.2g/l KH_2_PO_4_).

### DNA manipulation

Genomic DNA purification for use in cloning or whole-genome sequencing was purified using the Monarch HMW DNA Extraction Kit for Tissue (T3060L, New England Biolabs) according to manufacturer’s instruction with one modification: 1 ml overnight culture was washed with 1 ml PBS prior to first step. Plasmid DNA was purified using the Macherey Nagel Plasmid purification kit, according to manufacturer’s specifications. For cloning, DNA was routinely purified with the NucleoSpin gel and PCR clean-up kit (Macherey-Nagel) according to the manufacturer’s instruction or by ethanol precipitation, and verified by gel electrophoresis. DNA was quantified by gel electrophoresis or with a Qubit fluorometer (Thermofisher).

### Allelic replacement for gene deletion and genomic integration

Gene deletions or integrations were carried out according to Hmelo et al (36) using the allelic replacement vector pNJ1 (32). Briefly for construction of gene deletions, regions up- and downstream of target gene(s) were PCR amplified and joined by SOE PCR (specifics about methods can be found in Supplementary Methods and Supplementary table S 3). For deletion of the OSA cluster, the deletion construct fragments were 5.6, 5.6, and 5.3 kb for PA14, ST111 and PAO1, respectively. Constructs were ligated to pNJ1 after restriction digestion and used for transformation of calcium competent *E. coli*. These were then transferred to *P. aeruginosa* by conjugation by addition of helper strain containing pRK600. Briefly for genomic integration, a target region was PCR amplified and ligated to pNJ1 after restriction digestion.

### Direct cloning of serotype clusters in *P. aeruginosa*

RecET mediated recombineering was carried out according to Wang et al (26) using the pR6kT-mini-Tn7T-Km (31) vector to capture and transfer serotype gene clusters to *P. aeruginosa* with the following modifications: the donor DNA was tagged by integration of pNJ1 in the serotype cluster, to provide a selection for successful genomic capture. A detailed description of the individual steps can be found in Supplementary Methods and primers in Supplementary table S 3. Transfer of constructs to attTn*7* in *P. aeruginosa* was done according to Choi et al (31) by four-parental mating and verified by PCR. Selection markers, including the pNJ1 plasmid, were removed by sucrose counter-selection and subsequent flp/frt recombination.

### Direct cloning of fluorescent markers to mini-ctx2

PCR fragments from pBG42 (34) and pBG42 mKate were designed with overhangs homologous to mini-ctx2 (35) using PCR. These fragments contain *aacC1* (gentamycin resistance) and msfGFP or mKate. Plasmid preparation of mini-ctx2 was digested using SacI (Thermofisher) and purified. RecET recombineering was used to construct mini-ctx2 *aacC1* msfGFP and mini-ctx2 *aacC1* mKate. These plasmids integrate into the CTX attachment site in the *P. aeruginosa* genome and thus mediate genomic tagging of *P. aeruginosa* with *aacC1* and a green or red fluorescent marker. The plasmids were introduced into *P. aeruginosa* by conjugation according to Hoang et al (35).

### Whole-genome sequencing and assembly

Illumina sequencing was carried out on the Novaseq 6000 platform using 2×150 bp paired-end reads. Nanopore sequencing was barcoded using the rapid barcoding kit (SQK-RBK004) and sequenced on a R9.4.1 flowcell using the MinION device. Nanopore reads were basecalled using MinKNOW (3.1.19). FastP (0.12.4) (37) was using for quality control of Illumina reads, including adapter trimming and base correction (flag –c) for paired-end data. Reads were subset by seqtk (1.3-r106) (38) with random seeds for paired-end Illumina reads. For long-read sequencing, porechop (0.2.4) (39) was used for read demultiplexing and adapter trimming and reads were subset using filtlong (0.2.0) (40) with –min_length 1000. Hybrid and Illumina-only assemblies were performed using Unicycler (0.5.0) (41) and these were annotated by the prokaryotic genome annotation pipeline (PGAP) (v6.5) (42).

### Adhesion of *P. aeruginosa* strains

Bacterial biofilm formation in tissue culture plates was quantified according to protocols of O’Toole & Kolter (43) using crystal violet staining, to focus on the initial attachment of bacteria. Briefly, overnight cultures, grown in LB at 37 for up to 18 hours, were washed 2x in PBS by centrifugation at 5000*g* for 3 min. The OD_600_ of each culture was adjusted to OD_600_ = 1 in PBS and 200 µl culture was transferred to wells of a microtiter plate. 200 µl PBS was added to empty wells and used as blanks. Microtiter plates were incubated for 2 hours at 37°C with lid on, then washed 3x by flicking out liquid, and adding 250 µl distilled water to each well. Biofilms were stained by addition of 200 µl 0.1% crystal violet (freshly prepared on each day, dissolved in distilled water) for 15 minutes at room temperature. Plates were then washed 4x by flicking out liquid and adding 250 µl distilled water. 200 µl 96% ethanol was added to each well and the plate was incubated at room temperature for 30 minutes before measuring the 570nm absorbance of each well with a Biotek Cytation 5 or Synergy H1 plate-reader. Data was processed by an in-house script where the absorbance of blanks was subtracted from all absorbance values.

### Simulated partition coefficient of serotype repeat units

Chemical structures of the serotype repeat-units was characterized using ACD/ChemSketch 2014 using the addon logP (v.14.03) (44). Briefly, the known structures of the serotype repeat units (O4, O5, O12, O19) were re-created in ChemSketch according to (12, 17). Using the logP addon, the relative polarity (logP) of each serotype structure could be estimated.

### Lysis rate of bacteria in EDTA+Lysozyme solution

*P. aeruginosa* strains were lysed in EDTA and lysozyme according to Ayres et al (45). Briefly, bacterial cultures were grown overnight for 16-18 hours in LB broth at 37°C, 200 rpm. 1.5 ml culture was transferred to an Eppendorf tube and washed 3x by centrifugation at 5000*g* for 3 minutes in 1 ml 20 mM tris buffer (pH 7.8). The OD and volume of each culture was then adjusted to 1.2 ml OD = 1.25 in 20 mM tris buffer. A dilution row of EDTA was prepared in a microtiter plate using a multi-channel pipette. Lysozyme and bacterial suspension was added for a final concentration of 10 µg/ml and OD = 0.5, respectively, with a total volume of 250 µl in each well. After addition of bacterial suspension, the absorbance at 600nm was continuously read (at the lowest possible interval) with a Biotek Cytation 5 or Synergy H1 plate-reader at 30°C for 1 hour. The influence of magnesium was assayed in a similar way, but with a final concentration of 10mM MgSO_4_ in each well. The lysis rate in each well was determined from the kinetic curves using an in-house R script.

### Twitching motility

Twitching motility of *P. aeruginosa* strains was assayed according to Turnbull et al (46) on LB (1% Agar) plates. Briefly, a single colony was carefully touched with a 10µl pipette tip and stabbed through the agar layer to inoculate the petri dish plastic. Plates were inverted and stacked in a plastic box (3 plates/stack) and incubated at 37°C with the lid slightly ajar for 24 hours. An open petri dish containing a wet paper towel was placed inside the plastic box for humidity. Two perpendicular diameters were measured to quantify twitching motility of each assay, which was used to calculate the twitching motility area (*A* = *R*1 ∗ *R*2 ∗ *π*). To photograph twitching motility, the agar layer was carefully removed and the petri dish was photographed with a T:Genius gel imager using the UV transilluminator.

### Single cell atomic force spectroscopy

For single cell force spectroscopy experiments, bacterial probes were brought into contact with glass or polystyrene substrates (47, 48). Briefly, bacterial cultures were grown overnight for 16-18 hours in LB broth at 37°C, 200 rpm. These were washed twice in PBS by centrifugation and diluted 100-fold. Bacterial probes were subsequently prepared as previously described (49) by attaching single bacterial cells to hydrophobic AFM colloidal probes. AFM experiments were carried out in PBS at room temperature, using a JPK NanoWizard® 4 NanoScience instrument. Force-distances curves were recorded while approaching and retracting bacterial probes to and from the substrates on areas of 10 µm x 10 µm, using a constant approach and retraction speed of 3 µm/s, a ramp length of 3 µm and an applied force setpoint of 0.25 nN, at a resolution of 16 x 16 pixels. The adhesion properties of each bacterial strain were characterized by measuring the magnitude and frequency of force plateaus in the retraction curves (47, 50).

### Murine infection models

All animal procedures were conducted according to the guidelines of the Emory University Institutional Animal Care and Use Committee (IACUC), under approved protocol number PROTO 201700441. The study was carried out in strict accordance with established guidelines and policies at Emory University School of Medicine, and recommendations in the Guide for Care and Use of Laboratory Animals, as well as local, state, and federal laws. Mouse pneumonia infection was performed with six -to eight-week-old female BALB/c mice (Jackson Laboratories, Bar Harbor, ME) as previously described with a few modifications (51). Briefly, mice were anesthetized by intraperitoneal injection of 0.2 ml of a cocktail of ketamine (10mg/ml) and xylazine (5 mg/ml). *P. aeruginosa* strains: PAO1, IATS O4, IATS O12, ST111, ST111-2, PAO1ΔO+O4 and PAO1ΔO+O12 were grown on Difco^TM^ Pseudomonas Isolation Agar (PIA) for 16-18 hours at 37°C and suspended in PBS to an O.D_600_ of 0.5, corresponding to ∼10^9^ CFU/ml and inocula were adjusted to obtain the desired challenge dose.

For acute pneumonia model, mice were intranasally instilled with indicates *P. aeruginosa* doses in 25 μl of PBS. For systemic infection, mice were infected by intraperitoneal injection of desired dose in 0.1 ml of P. aeruginosa strains as indicated. To assess the bacterial burden, mice were euthanized at 24 h post-infection and whole lungs were collected aseptically, weighed, and homogenized in 1 ml of PBS. Tissue homogenates were serially diluted and plated on PIA for CFU enumeration. For survival studies, infected mice were monitored up to 3 weeks post-infection. Animals that succumbed to infection or appeared to be under acute distress were humanely euthanized and were included in the experiment results.

### LPS purification & visualization

LPS was purified from 1 ml *P. aeruginosa* overnight cultures grown on LB agar that were normalized to OD_600_ = 0.5 (∼1e9 CFU / ml). Cultures were pelleted by centrifugation and resuspended in 200 µl SDS sample buffer (BioRad) and briefly boiled. Samples were incubated with DNaseI (240 µg/ml), RNase (240 µg/ml) for 30 minutes at 37°C, followed by 3 hours incubation with Proteinase K (465 µg/ml) at 59°C. 10 µl purified LPS was loaded and separated on a 4-12% gradient Tris-glycine (BioRad) using SDS-PAGE and visualized by Western blotting or with the Pro-Q emerald lipopolysaccharide stain kit (Thermo Fischer) according to the manufacturer’s specifications. *P. aeruginosa* serogroup specific (O4, O12, O10/O19) rabbit polyclonal antibodies (Denka Seiken, Tokyo Japan) or *P. aeruginosa* anti O5 mouse IgM MF15-4 monoclonal antibody (MediMabs, cat # MMM-76605-1) were used for Western blotting. Anti-mouse IgM-HRP or anti-rabbit IgG-HRP were used as secondary antibodies, respectively.

### Serum bactericidal assay

Overnight cultures were diluted in PBS supplemented with 1% proteose peptone to OD_600_ = 0.05 for a final inoculum of approximately 1e6 CFU per well. Normal human serum (NHS) was diluted in PBS plus 1% proteose peptone to twice the final concentration. As controls, cultures were additionally incubated with NHS that was heat inactivated (HI) by incubation at 56°C for 30 minutes, or PBS +1% proteose peptone (0% serum). Equal volumes (50 µl) of sera and bacterial suspensions were mixed and incubated at 37°C for 1 hour with gentle shaking. An aliquot from each well was serially diluted and plated on LB agar for CFU enumeration after incubation overnight at 37°C. Serum survival was determined for each strain by determining the log10 fold difference between CFU recovered after incubation with NHS and HI.

### Statistical tests and data analysis

Rstudio (2022.07.2 build 576) was used for statistical testing. For comparison of multiple means, the compare_means function of ggpubr (52) was used with method = “t.test” and correction for multiple testing with false discovery rate (FDR). For comparisons of means between two means, the built-in function t.test from R stats was used (Welch’s t-test). The results of animal survival studies are presented using Kaplan-Meier survival curves. The Logrank test (survdiff) of package survival (53) was used to determine if survival curves were different. Other R Packages used for data analysis and visualization: ggplot2 (54), readr (55), viridis (56), and ggfortify (57). Significance level (p value) of statistical tests is indicated by ns for non-significant, * for p<0.05, ** for p<0.01, *** for p<0.001, and **** for p<0.0001.

## Results

### Synthetic serotype switching by genomic capture and tn7 integration

To investigate the impact of serotype switching in the evolution of HIRIC ST111, we designed a method that enabled the construction of isogenic strains each expressing different O-antigens (i.e. synthetic serotype switching). This method includes genetic tagging of the O-antigen (OSA) gene cluster with an antibiotic resistance gene to facilitate genomic capture, preparing a mini-tn7 vector for recombination, and RecET recombineering for the construction of a serotype switch plasmid as illustrated in Figure 1ab. The method enables the capture of the entire O-antigen cluster from a donor strain of choice, and site-specific integration at attTn7 in *P. aeruginosa* (Figure 1c) followed by removal of selection markers and plasmid constructs. Synthetic serotype switching is then accomplished by first deleting the native OSA cluster (16.6 kb in PA14, 22.5 kb in PAO1, and 14.3 kb in ST111) using allelic replacement (Materials and Methods) and then integrating a heterologous O-antigen cluster by tn*7* tagging. This approach was used to construct isogenic strains expressing serotypes O4 or O12 in the well-characterized strains PAO1 (serotype O5), PA14 (serotype O19) and a serotype O4 isolate of ST111, to understand the biological effect of the O4-to-O12 serotype switch in different genetic backgrounds (Table 1).

**Figure 1.**
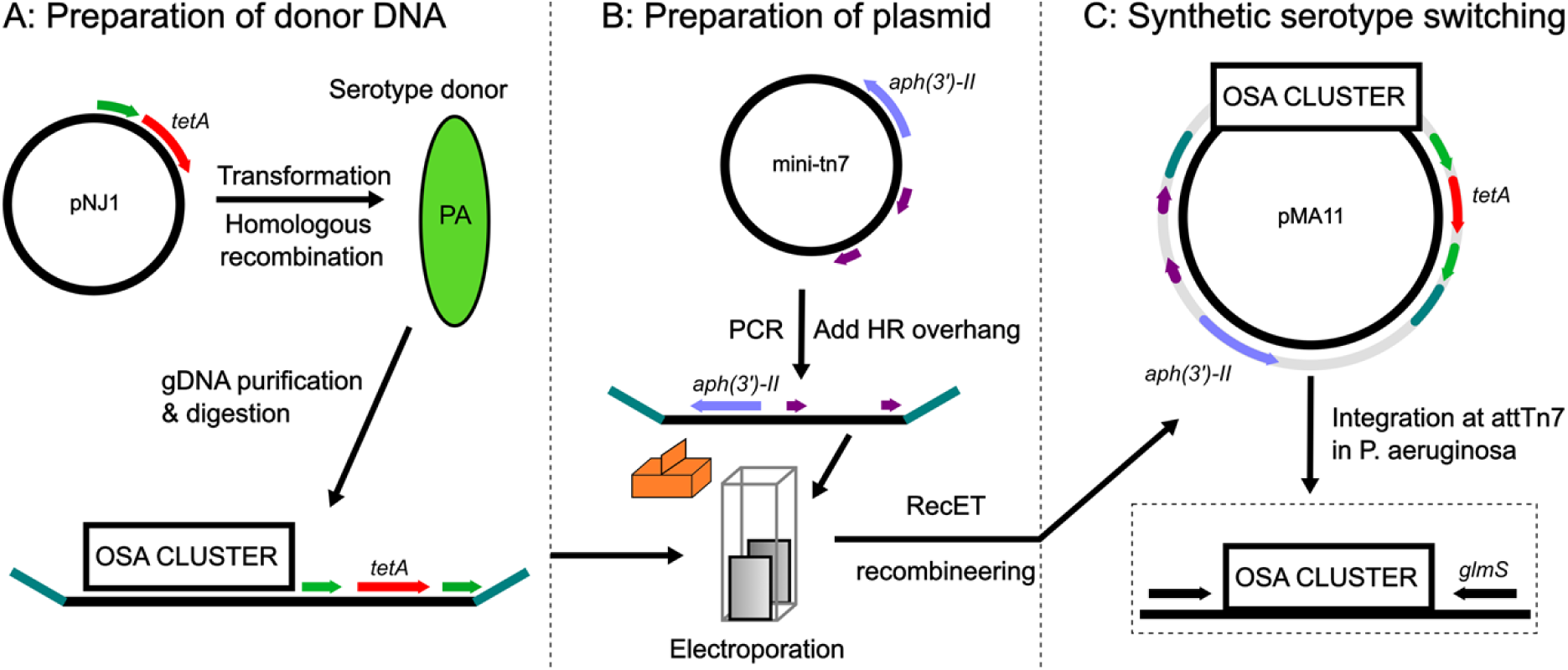
Synthetic serotype switching in *P. aeruginosa* by RecET genomic capture and tn7 tagging. A: Donor serotype strains are prepared by integration of pNJ1 in the serotype cluster by homologous recombination. This vector (pNJ1) contains a 900 bp PCR fragment homologous to the serotype cluster (green), *tetA (red),* and *sacB* (not pictured). Genomic DNA is purified and digested by restriction enzymes that cut up- and downstream of the serotype cluster (within 5000 bp). B: Genomic tagging plasmid (pUC18R6kT-mini-Tn7T-Km) is linearized by PCR to add 80 bp overhangs (teal) which are homologous to regions up- and downstream of serotype clusters. Serotype donor DNA and linearized plasmid are mixed and electroporated into competent Gbdir-*pir* cells which facilitate RecET recombineering. C: The resulting plasmid can facilitate synthetic serotype switches by integration at attTn7 in *P. aeruginosa*, which is upstream of *glmS*. DNA between Tn7L and Tn7R (purple) is integrated at attTn7 in the presence of helper plasmid pTNS2. Antibiotic genes *tetA* and *aph(3’)-II* were excised by sucrose counter selection and FLP recombination, respectively.

### Verification of strains expressing heterologous O-antigens

To verify that our approach resulted in isogenic strains expressing heterologous O-antigens, we first purified and separated LPS on SDS PAGE gels and visualized this with Pro-Q Emerald 300 LPS stain or Western blotting, using antibodies specific for different O-antigen serotypes (Materials and methods). We found that all strains reacted with the LPS stain (Figure 2a-b) and the visualized O-antigen LPS banding indicates functional heterologous expression of OSA in all engineered strains. As expected, no LPS bands were observed in strains without an OSA gene cluster. These results were further confirmed with Western blot analyses using corresponding serotype specific antibodies (Figure 2c-f). Interestingly, it was evident that the OSA length regulation is dependent on the genetic background of the strains used for heterologous expression. For example, as seen in Figure 2ab, the LPS size distribution is different among most of the strains. In IATS O12 the “very long” OSA chains seems to be the predominant LPS glycoform, whereas in ST111, PA14, PAO1 there is a mix of “long” and “very long” OSA chains. This glycoform diversity between genetic backgrounds is to be expected as the gene that regulates the “very long” OSA chains resides outside the OSA locus (58). Lastly, genome sequencing of the engineered strains confirmed the integrity of OSA gene cluster and that no genetic changes had been introduced to the heterologous OSA cluster during strain construction (Supplementary Table S1).

**Figure 2.**
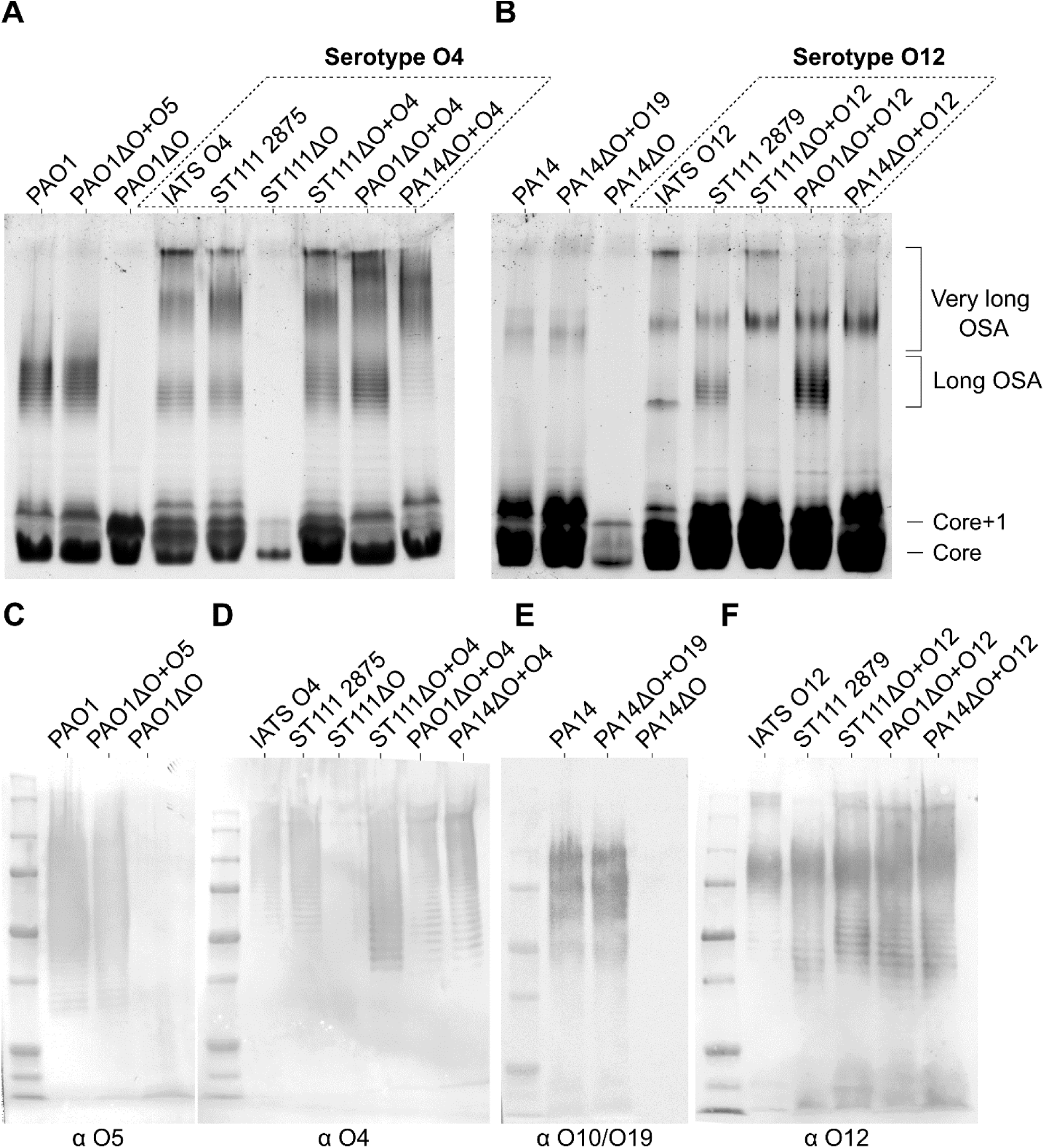
Pro-Q and Western blot LPS visualization of P. aeruginosa strains. A & B: LPS separated by electrophoresis on SDS PAGE 4-20% gels and stained with Pro-Q, strain names have been shown above their corresponding lanes. The location of different LPS glycoforms have been indicated on the gels. C-F: Western blot gels using antibodies specific for P. aeruginosa serogroups (primary antibody shown below each Western blot membrane). C: MF15-4 Mouse-IgM (monoclonal), D: Rabbit anti sero O4 (polyclonal), E: Rabbit anti sero O10/O19 (polyclonal), F: Rabbit anti sero O12 (polyclonal). For Western blots, anti-mouse IgM-HRP or anti-rabbit IgG-HRP were used as secondary antibodies, as appropriate.

### Serotype O12 adheres better to tissue culture polystyrene than O4

We first investigated if expression of different O-antigens affects the ability of the strains to adhere to tissue culture polystyrene microtiter plates. To do this, we used a variant of the classical crystal violet assay (43) wherein washed cell suspensions were incubated for 2 hours in microtiter plates without nutrients. After removal of planktonic and weakly attached cells, the adhering cells are quantified by crystal violet staining.

Using this assay, we find that the three wild-type strains (PA14, PAO1, and ST111) exhibited different levels of adhesion (Figure 3), and that deletion of the OSA gene cluster resulted in a significant decrease in adhesion for all strains (cf. wildtype and ΔO (OSA deletion strains) strains in Figure 3). For PA14 and PAO1, we find that heterologous expression of OSA restores adhesion, but that only expression of O12 in ST111 results in an adhesion phenotype distinguishable from the OSA deletion strain. The results obtained from ST111 suggest that heterologous expression of either O12 or O4 could not complement the adhesion phenotype of the OSA deletion strain to reach wild type (O4) levels. This was not due to lack of OSA production (Figure 2), and we speculate that this phenotype is a result of mutations that inactivate other (non-OSA) adhesion factors.

**Figure 3.**
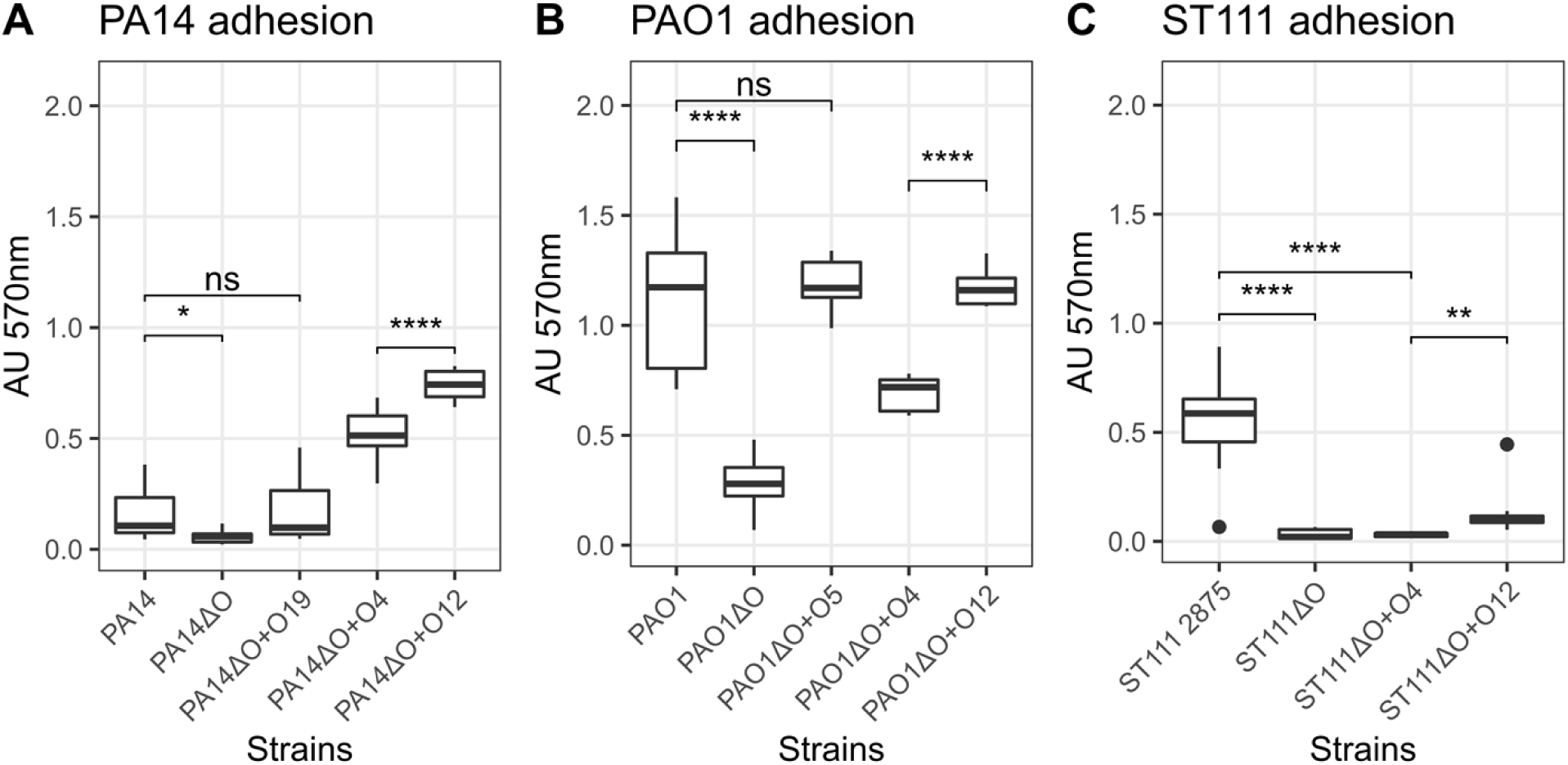
O-antigen structure changes adhesion of strains in PA14 (A), PAO1 (B), and ST111 (C). The adhesion of strains (shown on y-axis, common to all plots) was measured after 2 hours by crystal violet staining. All strains were characterized using 12 biological replicates and 6 technical replicates. Common for all strains is that loss of the O-antigen severely impairs the ability of the bacteria to adhere to tissue culture polystyrene plates (p>0.05), and complementation with the wild-type serotype restores adhesion in PA14 (O19) and PAO1 (O5). Surprisingly this is not the case for ST111 (O4) where all of the engineered strains show impaired adhesion compared to the wild type. For all strains tested we find that serotype O12 strains are significantly more adherent than serotype O4 strains (p>0.05).

Importantly, in all strain backgrounds we find that heterologous expression of O12 is associated with significantly higher adhesion than O4 expression. Specifically, in the PA14 background we find that serotype O12 is 1.4x more adherent than serotype O4 (*p*=2.2e-5, multiple t-test w/ fdr) and that serotype O12 is 1.7x more adherent than the O4 strain (*p*=5e-12, multiple t-test w/ fdr) in PAO1. Despite the lowered adhesion capabilities of all engineered ST111 strains, we nevertheless find that O12 is 4.3x more adherent than the engineered O4 strain (*p*=9.2e-3, multiple t-test w/ fdr)

Based on the results of these experiments we conclude that the molecular structure of the O-antigen has a significant effect on the early stages of biofilm formation (adhesion) as demonstrated by attachment to tissue culture polystyrene, and that serotype O12 facilitates better adhesion than O4 regardless of the genomic background.

### Relative polarity of OSA structures plays a role in bacterial adhesion

Bacterial adhesion is governed by many physiochemical forces including polarity (59–61). To test the influence of polarity on the adhesion of strains to tissue culture polystyrene, we modelled the relative polarity (Partition coefficient logP = log [A]_org_-log [A]_aq_ , see materials and methods) of a single OSA repeat unit for each serotype explored in this study (O4, O12 as well as parental serotypes O5 and O19 in PAO1 and PA14, respectively). In terms of relative predicted hydrophilicity, the serotypes rank O5 > O12 > O4 > O19. We find a significant association between the relative polarity of OSA structures (exponent of logP) and the experimental adhesion data of both PAO1 and PA14 (absorbance at 570nm) using linear regression models (p<0.05, t-test) (Figure 4a). While this model is not sufficient to explain all variance (R^2^ shown in Figure 4a), it shows that polarity of the OSA does play a substantial role in adhesion to tissue culture polystyrene, in a strain specific manner as PAO1 strains can be seen adhering at a higher level than PA14 strains despite expressing the same serotypes.

**Figure 4.**
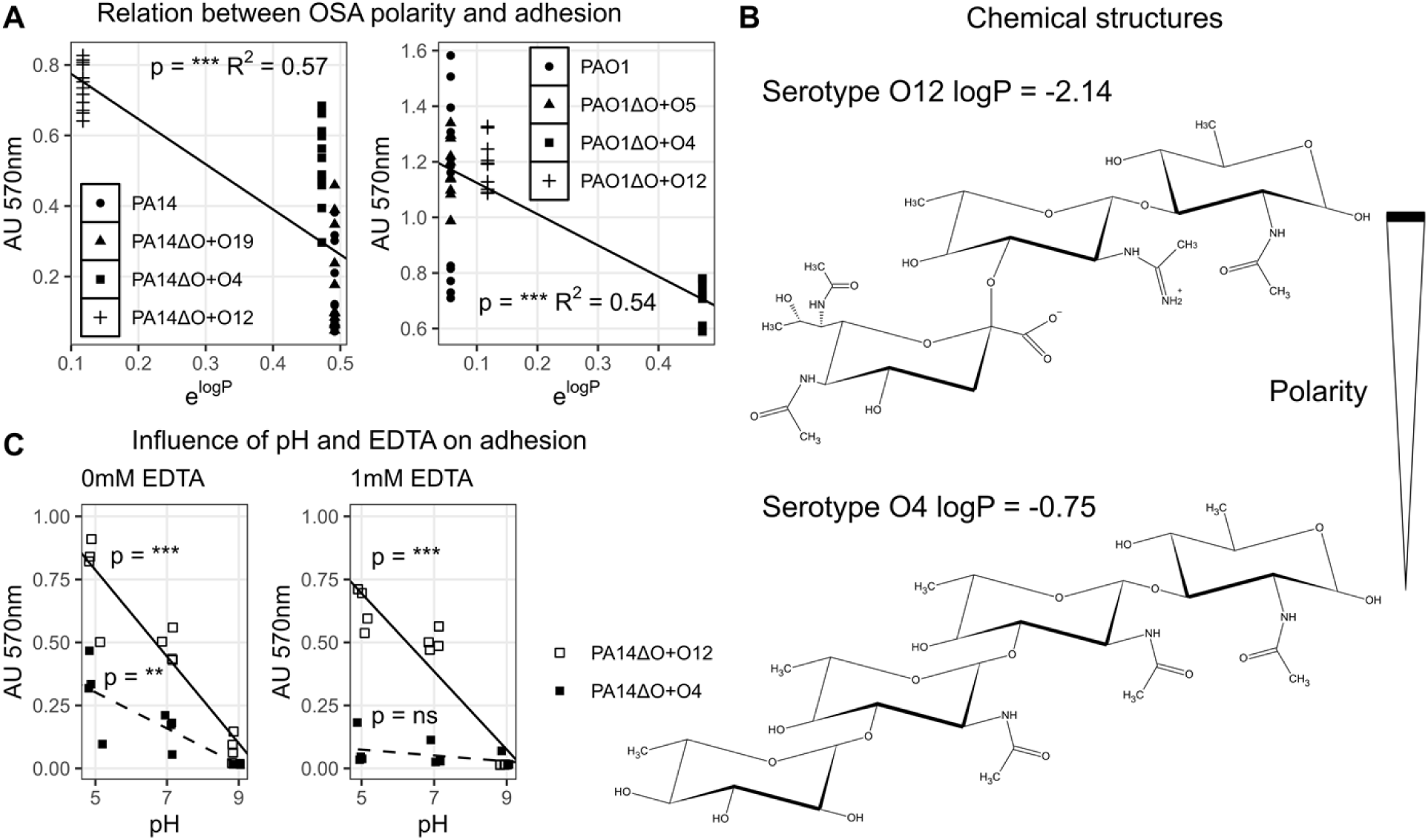
Relating the biophysical forces that govern bacterial adhesion to experimental data. A: We show that the relative polarity of the expressed serotype correlates with adhesion to tissue culture polystyrene. This is the case for PA14 (left) and PAO1 (right) strains expressing different serotypes. Significant (p and R^2^ values shown in each plot) linear models between adhesion and the polarity of OSA repeat units are made for each strain and shown as lines between the data points. B: The chemical structure of the OSA repeats for serotype O12 (top) and serotype O4 (bottom) along with their relative polarity (logP). In terms of relative polarity, these figures indicate that the O12 repeat unit is more polar than the O4 repeat unit. C: The influence of pH and EDTA on adhesion. The adhesion of PA14ΔO+O4 and PA14ΔO+O12 was assayed at pH 5, 7, and 9 in the absence (left) and presence (right) of the metal chelator 1mM EDTA. Linear models of adhesion as a function of pH were made for both strains and shown as a line, for PA14ΔO+O12, or a dashed line, for PA14ΔO+O4. Significance of linear models between adhesion and pH is shown in the figure above their corresponding lines. In the absence of EDTA, adhesion of both strains increases as pH decreases. The electrostatic repulsion between the negatively charged substrate and bacteria is reduced, as negatively charged phosphates in the LPS become increasingly protonated at decreasing pH. In the presence of EDTA, adhesion of PA14ΔO+O4 is abolished, while PA14ΔO+O12 is still able to adhere, suggesting that serotype composition plays a major role in binding LPS stabilizing divalent cations.

### Serotype O12 maintains adhesion in the presence of EDTA, whereas O4 does not

Another key aspect of bacterial adhesion is the influence of electrostatic interactions, which normally represent repulsive forces between bacteria and substrate (tissue culture polystyrene) since both are negatively charged (59, 62–64). The surface charge of *P. aeruginosa* is predominantly mediated by phosphate groups in the LPS core, which serve to increase surface electronegativity. When the LPS is decorated with OSA, the overall surface electronegativity is reduced by divalent cations, which stabilize the outer membrane (13, 45, 59, 65, 66). Due to the different chemical properties of serotypes O4 and O12 (Figure 4b), we hypothesize that the adhesive properties of each serotype are differentially affected by pH and the presence of the metal chelator EDTA. To test this, we assayed adhesion of isogenic strains expressing either O12 or O4 (PA14ΔO+O4 and PA14ΔO+O12) at pH 5, 7, and 9 in the absence or presence of 1mM EDTA.

We find a significant linear correlation between pH and adhesion for both serotypes O12 and O4 (significance levels indicated in both panels of Figure 4c), showing that adhesion increases as pH decreases. We assume that the physiochemical basis for increased adhesion with decreasing pH is due a reduction in the charge of phosphate groups in the LPS (67), which in turn decreases the repulsion between the negatively charged bacteria and tissue culture polystyrene. However, in comparing the slopes for each serotype, we see that the effect of pH on adhesion is 2.4 times greater for serotype O12 (slope: -0.17) than O4 (slope: -0.07) (Figure 4c, left panel). We propose that this difference is in part due to the positively charged acetamidino group in the OSA of serotype O12 (68, 69) (Figure 4b) and in part due to the ability of serotype O12 to mask its surface charge by binding more divalent cations with the carboxyl group (59) than serotype O4.

Importantly, adhesion of PA14ΔO+O4 is greatly reduced upon addition of EDTA (Figure 4c, right panel), whereas adhesion of PA14ΔO+O12 is unaffected by EDTA. We propose that the same functional groups in the OSA that differentiate these strains response to pH (acetamidino and carboxyl), can maintain adhesion in the presence of EDTA. Overall, these results show that chemical differences between O12 and O4 impact adhesion behaviors of cells as well as the sensitivity of these behaviors to changing environmental conditions.

### Cell physiology and motility behavior is modulated by the O-antigen composition

Since our results suggest that O4 and O12 O-antigens have different capacities to bind LPS stabilizing cations (such as Ca^2+^ and Mg^2+^), we hypothesized expression of different O-antigens would affect membrane stability and consequently also the functionality of membrane associated proteins. To test this hypothesis, we first investigated if EDTA exposure would result in differential increase in outer membrane permeability in cells expressing either O4 or O12 O-antigens. *P. aeruginosa* is known to be highly resistant to lysozyme, but becomes lysozyme-sensitive when co-treated with EDTA which makes the outer membrane of *P. aeruginosa* permeable (45). Measurements of the lysis rate of cultures exposed to 1 mM EDTA and 10 mg/mL lysozyme revealed that PA14ΔO+O12 is less sensitive to the treatment than PA14ΔO+O4, suggesting that the membrane stability is influenced by the chemical composition of the O-antigen (Figure 5a). We additionally show that addition of magnesium restores membrane stability (and thus resistance to EDTA+lysozyme treatment), which confirms the role of divalent ions in this interaction.

**Figure 5.**
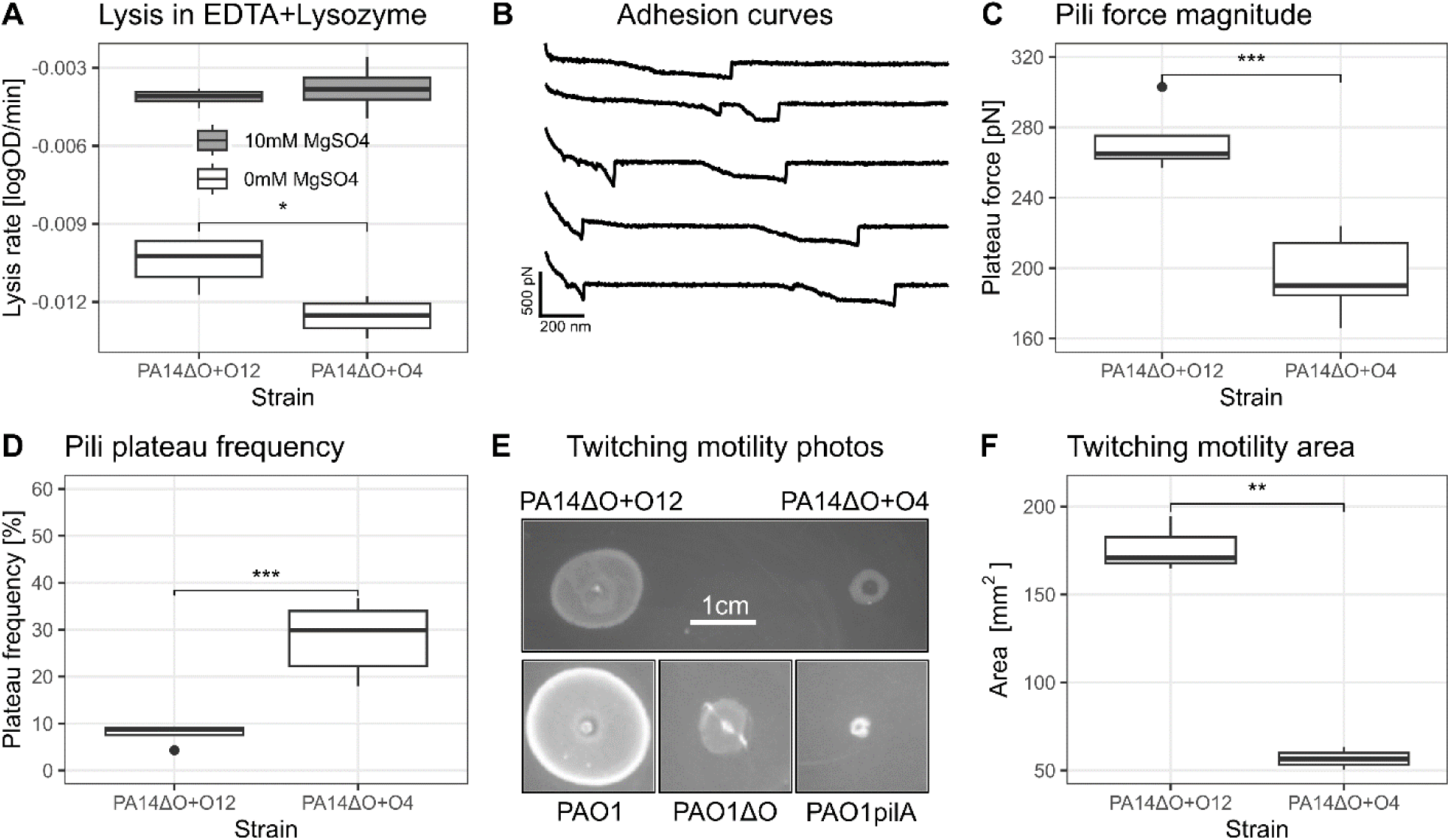
OSA composition plays a role in a multitude of bacterial interactions not directly related to LPS. A: The serotype significantly affects the rate of lysis when treated with 1 mM EDTA & lysozyme. This effect can be rescued by the addition of magnesium, implicating the role of divalent cations in this interaction. Each boxplot represents data from 4 biological replicates. B-D: Atomic force microscopic (AFM) characterization of serotype switched strains shows that pili mechanical properties are significantly affected by serotype composition; B: Representative adhesion curves for PA14ΔO+O4 from which we can identify type IV pili by their characteristic plateau signatures. C: Strength and D: frequency of force plateaus for PA14ΔO+O12 and PA14ΔO+O4 strains. E: Serotype composition affects twitching motility as can be seen comparing the motility of PA14ΔO+O12 and PA14ΔO+O4 (grown on the same plate, inoculated simultaneously). Prior understanding about twitching motility is that the O-antigen is essential for twitching motility (73), but here we can show that this is not the case with isogenic PAO1 strains deficient in either O-antigen (PAO1ΔO) or type IV pili (PAO1pilA). Scale bar (1 cm) is common to all twitching motility photos. F: The twitching motility area is significantly affected by serotype composition.

If membrane stability is affected by the chemical composition of O-antigen, it could be expected that the function of proteins or protein complexes associated with the membrane would also be affected by different O-antigen structures. To explore this idea, we focused on Type IV pili (T4P) and used atomic force microscopy (AFM) to investigate at single-cell level, the mechanical properties of T4P in cells expressing different O-antigens.

To quantify the forces between *P. aeruginosa* polystyrene substrates, we used single cell force spectroscopy (SCFS) (48, 70). A single bacterial cell is attached to an AFM colloidal probe and brought into contact with a polystyrene surface, while recording force-distance curves of the approach and retraction of the AFM cantilever. The adhesive interactions observed in the retraction curves are established through interactions between biological polymers (proteins, polysaccharides) at the surface of the bacterium and the polystyrene surface. The features observed in the adhesion curves are very often specific of certain types of biological structures, such as the type IV Pili force plateau signature (47, 50). When characterizing the adhesion of PA14 strains (PA14ΔO+O4 and PA14ΔO+O12) to polystyrene, we identified the relative contribution of type IV pili through force plateaus (figure 5b) and found a significant difference in the adhesion force of pili expressed by the PA14ΔO+O12 (284 ± 37 pN) strain compared to PA14ΔO+O4 (195 ± 39 pN) (Figure 5c). Interestingly, the frequency of force plateaus is lower for the PA14ΔO+O12 strain (8 % vs 28 %) (Figure 5d), which may indicate that type IV Pili expressed by the PA14ΔO+O12 strain are mechanically more stable than those expressed by the PA14ΔO+O4 strain. This result supports our hypothesis that the O-antigen composition plays a role in the function of membrane associated proteins (specifically the type IV pili) due to its role in stabilizing the outer membrane of the bacteria.

T4P are known for its essential role in twitching motility, which allows *P. aeruginosa* to move across solid or semisolid surfaces by the extension and retraction of pili (46, 71, 72). As we have shown that the serotype composition alters the biophysical function of pili, we next tested if O-antigen composition affected twitching motility phenotypes. We found that the twitching motility area of PA14ΔO+O4 is substantially reduced compared to PA14ΔO+O12 (Figure 5e & f). We note that twitching motility is not abolished in the O4 or OSA deficient strains as observed in a twitching motility deficient PAO1pilA mutant (Figure 5e) (additional twitching motility data shown in Supplementary Figure S1). Overall, these results confirm our hypothesis that the serotype composition plays a role in cell physiology and alters bacterial behaviors that depend on the function of membrane-associated protein complexes.

### Serotype composition plays a major role in a murine acute pneumonia infection model

Given the differential impact of O-antigen structures on adhesion phenotypes and membrane stability (which influences the sensitivity towards lysozyme and T4P-dependent motility phenotypes), we next investigated the impact of O-antigen structures on virulence. To address this, we assessed the virulence of isogenic serotype-switched PAO1 strains in a murine acute pneumonia infection model using BALB/c mice. We chose the PAO1 derived strains for this experiment as PAO1 is well studied in this infection model. Mice were intranasally infected with approximately 2.5e+7 CFU and either euthanized 24 hours post-infection to estimate lung colonization (Figure 6a), or monitored for survival up to 7 days post-infection (Figure 6b).

**Figure 6.**
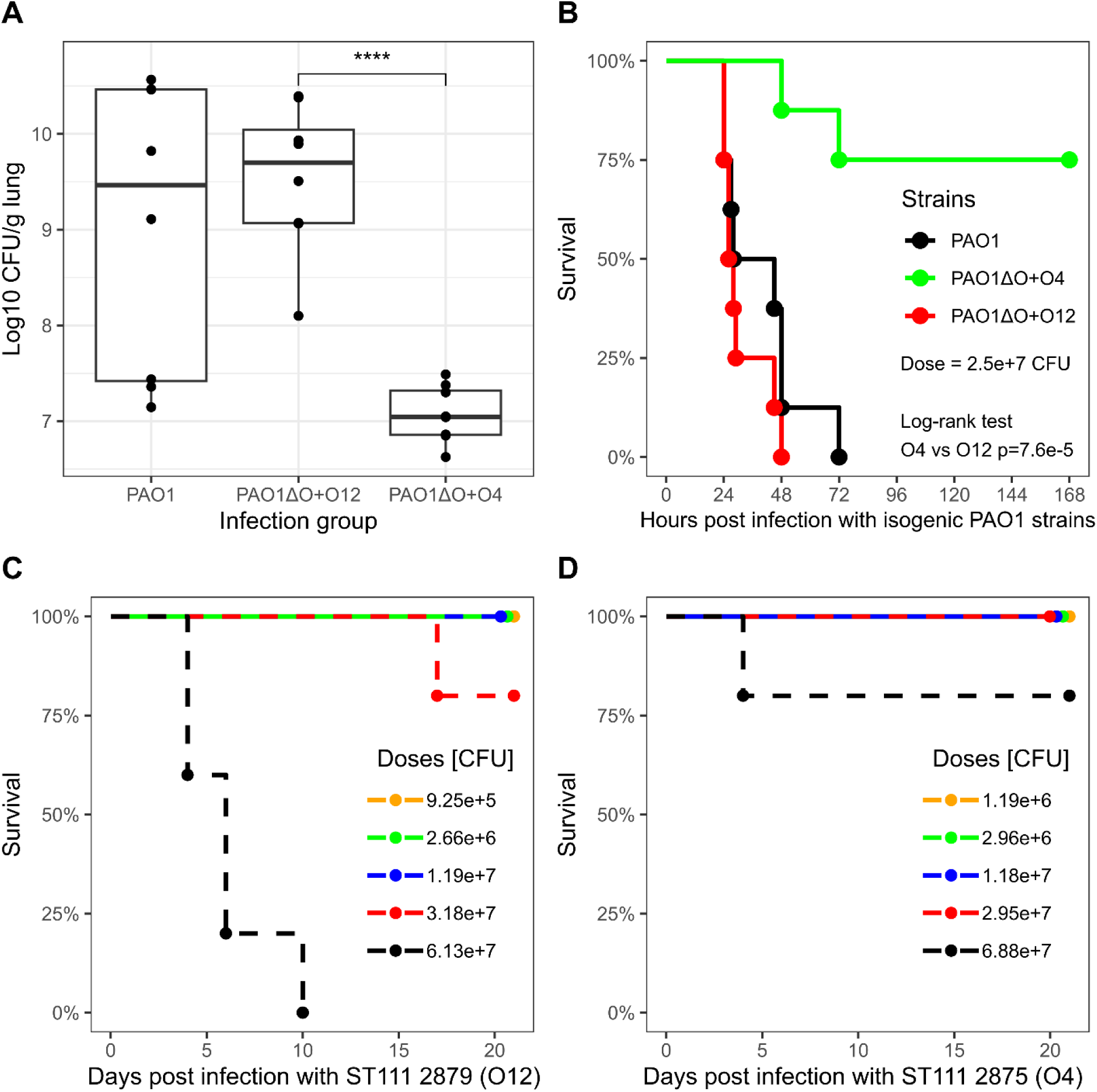
Virulence characterization in clonal and isogenic serotype strains of P. aeruginosa expressing different O-antigens. Groups of mice were infected intranasally with the indicated dose of bacteria in an acute pneumonia infection model using BALB/c mice. A: 24h post-infection (dose ∼ 2.5e+7 CFU), whole lungs were aseptically collected from euthanized animals (n=8) and homogenized. CFU was estimated from homogenates by plate counting on selective media (PIA). B: Mice infected (n=8) with isogenic PAO1 strains expressing different O-antigens were monitored for survival 7 days post infection. C & D: Multiple groups of mice (n=5) infected with a clinical strain ST111 2879 expressing serotype O12 (C) or ST111 2875 expressing serotype O4 (D) monitored for survival 20 days post infection.

To determine whether the serotype composition impact the infection and colonization outcome, we monitored the lung colonization 24 hours post-infection. Higher bacterial loads were recovered from mice infected with wild-type PAO1 and PAO1ΔO+O12 strains. In contrast, the bacterial burden was significantly lower in the lungs of mice infected with PAO1ΔO+O4 (Figure 6a). In comparing the virulence of PAO1, PAO1ΔO+O12, and PAO1ΔO+O4, we find that expression of serotype O4 is associated with less virulence compared to wild-type PAO1 and the O12 expressing strain. As shown in Figure 6b, mice infected with either wild-type PAO1 or PAO1ΔO+O12 succumbed to infection by 48-72 hours. In contrast, better survival (75%) was observed in mice infected with PAO1ΔO+O4 (Figure 6b). Both of these results indicate a substantial difference in virulence between expressing serotype O4 and O12.

As we could demonstrate a clear link between serotype composition and virulence in this infection model, we sought to investigate if this difference also exists in clinical ST111 strains expressing either O4 or O12. Although these strains are not isogenic (27), which makes it difficult to attribute virulence phenotypes to specific O-antigens, the results would nevertheless provide insight into why ST111 O12 is more successful than ST111 O4.

Multiple groups of mice were intranasally infected with a range of different infection doses using ST111 strains 2875 (expressing O4) and 2879 (expressing O12) (Materials and methods). Surprisingly, unlike PAO1 infections where mice develop acute pneumonia symptoms by 24-48 hours post infection, only mice infected with a high dose (approx. 6e7 CFU) of ST111 serotype O12 strain displayed delayed onset of symptoms and succumbed to infection starting day 5 post infection (Figure 5c). The difference in virulence in ST111 expressing either O12 or O4 appear to corroborate the observations from isogenic PAO1 strains, but we cannot rule out that other genetic differences between ST111 strains 2875 and 2879 may contribute to their dissimilar virulence phenotypes.

### Composition of the OSA may play a role in serum resistance

One model to explain why serotype O12 may be more virulent than serotype O4 in the acute pneumonia infection model is that these strains exhibit an increased survival within the host. To address this, we sought to characterize the inhibition by complement-mediated killing of isogenic strains expressing different serotypes when incubated with normal human serum (NHS). Previous literature has described the importance of O-antigen length and presence in this interaction (58, 74); however, it is unknown what role the specific O-antigen composition plays.

Strains were incubated in either NHS or heat-inactivated serum (HI), as a control, for 1 hour. After incubation, the survival of each strain was quantified by CFU enumeration. We found that there is an association between loss of OSA and reduced fitness in both PAO1 and ST111 (c.f. WT and ΔO in Table 3). Complementing the PAO1ΔO strain with any serotype (O4, O12, or O5) restored survival in serum to similar levels as the wild-type PAO1 (Table 3), indicating that the chemical composition of O-antigen does not differentially affect serum resistance.

**Table 3.**
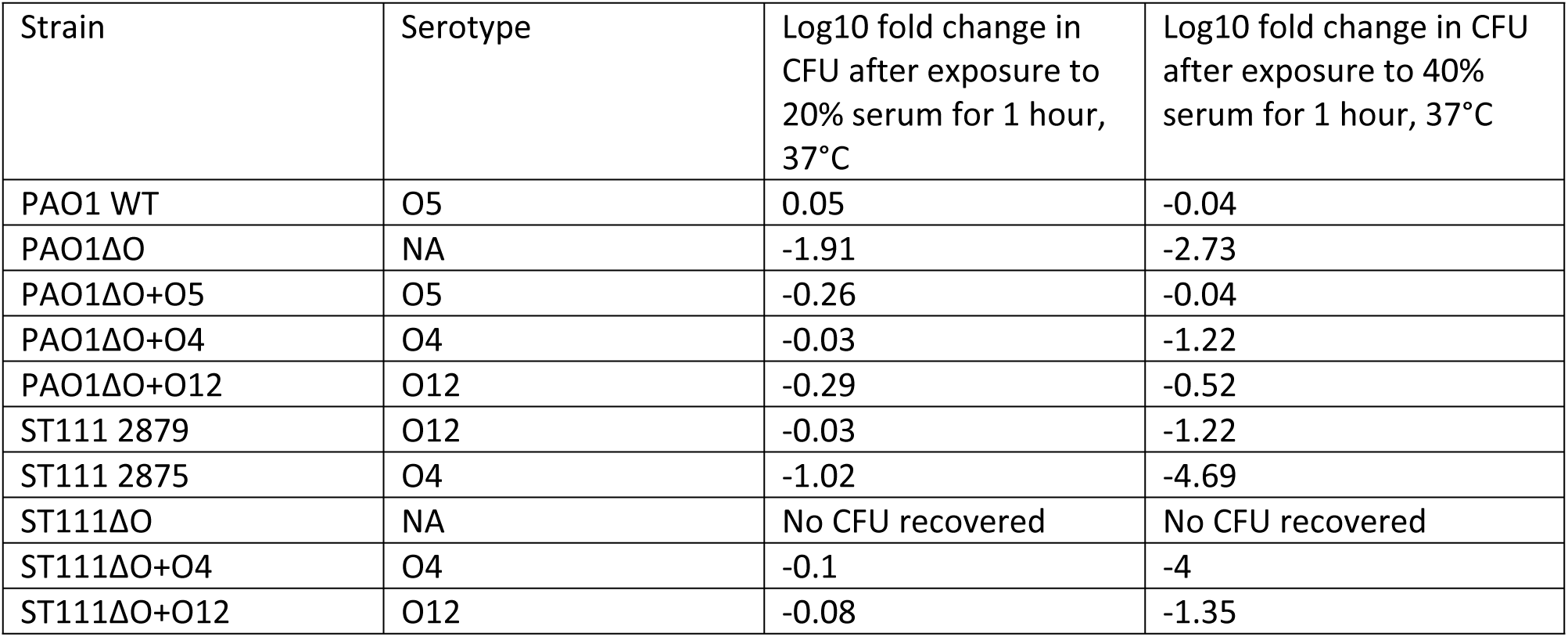
Survival of P. aeruginosa strains when incubated with normal human serum. Survival is quantified by determining the log10 fold difference between CFU recovered after 1h incubation in normal human serum and incubation in heat-inactivated serum (for 20% NHS n=4 and for 40% NHS n=3). NA: non-applicable; strains that do not express an O-antigen.

| Strain | Serotype | Log10 fold change in CFU after exposure to 20% serum for 1 hour, 37°C | Log10 fold change in CFU after exposure to 40% serum for 1 hour, 37°C |
| --- | --- | --- | --- |
| PAO1 WT | O5 | 0.05 | -0.04 |
| PAO1ΔO | NA | -1.91 | -2.73 |
| PAO1ΔO+O5 | O5 | -0.26 | -0.04 |
| PAO1ΔO+O4 | O4 | -0.03 | -1.22 |
| PAO1ΔO+O12 | O12 | -0.29 | -0.52 |
| ST111 2879 | O12 | -0.03 | -1.22 |
| ST111 2875 | O4 | -1.02 | -4.69 |
| ST111ΔO | NA | No CFU recovered | No CFU recovered |
| ST111ΔO+O4 | O4 | -0.1 | -4 |
| ST111ΔO+O12 | O12 | -0.08 | -1.35 |

At 20% NHS, there are only subtle differences between ST111 strains (cf. ST111 2875 and ST111 2879 in Table 3). However, at 40% NHS, these differences become more pronounced and the serotype O12 strain ST111 2879 is associated with better survival in serum than serotype O4 strain ST111 2875, which may directly correlate to the observed virulence difference between these strains. Similarly, ST111ΔO+O12 is associated with 2-4 logs increased survival compared to survival of ST111 2875 or ST111ΔO+O4, suggesting that O-antigen composition plays a role in serum resistance in this genetic background.

## Discussion

A limited number of MDR/XDR *P. aeruginosa* sequence types (e.g. ST111, ST244, and ST253) are highly prevalent in hospitals worldwide. To limit the continuing spread of these clones and to predict the next waves of additional MDR/XDR *P. aeruginosa* high-risk clones, it is important to determine the mechanisms that enables these particular clones to succeed while others do not. In this study, we examined how different O-antigen serotypes modify *P. aeruginosa* pathophysiology, since serotype switching from O4 to O12 O-antigens has previously been shown to be a characteristic of the emergence and success of ST111 as a high-risk clone.

To pinpoint how expression of different serotypes affect *P. aeruginosa* biology, we engineered and analyzed isogenic strains each expressing either O4 or O12. We find that the sugar composition of O-antigens affect adhesion in both laboratory strains (PAO1 and PA14) and in clinical ST111 strains (Figure 3). Our results in relation to adhesion extend previous studies that have clearly demonstrated that the LPS and the O-antigen plays an important role in adhesion and the initial stages of biofilm formation by studying knockout mutants to the O-antigen and the core (59, 74–78). These studies have shown how the different LPS glycoforms of *P. aeruginosa* (OSA-capped, CPA-capped, uncapped) can mediate different interactions with biotic and abiotic surfaces. Here, we find that our O-antigen deficient mutant strains are the most impaired in adhesion to the hydrophilic (79) tissue culture polystyrene plates, and we expect these strains are only able to express the hydrophobic uncapped LPS glycoform, due to a lack of WbpL (17, 80–84), or a truncated capped core in the case of PA14 (85). This is in agreement with previous literature that find O-antigen deficient strains to be more hydrophobic (59, 74, 86). When an O-antigen gene cluster is introduced and expressed in these mutants, we find that the relative adhesion of the engineered strains to the polar tissue culture polystyrene substrate follows the ranking O12 > O4 > ΔOSA, regardless of strain genotype. We propose that this difference in adhesion is mediated, in part, by the polarity of the O-antigen repeat unit. In support of this idea, we find that there is a significant correlation between the predicted partition coefficients (relative measure of polarity, logP) of serotype repeat units and the results from adhesion experiments. Despite the limitations of this model (i.e. it only takes the contribution of a single OSA repeat unit into account, and it does not account for charged groups and many other important physiochemical forces that we know play a role in cell adhesion (60, 87)), the results suggest that *P. aeruginosa* adhesion properties may be predicted on the basis of the sugar composition of the O-antigen.

Importantly, we observed a serotype specific impairment of adhesion for serotype O4 when adding the metal chelator EDTA. EDTA is known to destabilize the outer membrane of *P. aeruginosa* by sequestering cationic metal ions from the cell envelope (13, 65) and we propose that the observed difference in EDTA susceptibility (on adhesion) is the result of distinct chemical properties of each OSA structure (i.e. O4 versus O12). We further substantiate this finding by showing that the rate of EDTA-mediated lysozyme lysis of these strains is different and that serotype O4 is associated with a significantly higher rate of lysozyme lysis than serotype O12. The chemical differences in OSA composition that we propose are responsible for this difference include the number of ionizable groups (20, 68, 88), polarity (as discussed in the previous paragraph), and capacity to bind LPS stabilizing cations (such as Ca^2+^ and Mg^2+^), which can have an effect on membrane stability (14, 17, 89).

The finding that outer membrane stability is affected by the chemical composition of O-antigen suggests that expression of O4 and O12 O-antigen would differentially impact the function of proteins or protein complexes that are sensitive to outer membrane structure and function. Indeed, using single-cell atomic force microscopy (AFM), we show that the strength (Figure 5c) and frequency (Figure 5d) of Type IV pili (T4P) is different in O4 expressing cells compared to O12. These differences in biophysical properties of T4P also manifested in reduced twitching motility of O4-expressing cells (Figure 5ef). A link between O-antigen biosynthesis and twitching motility in *P. aeruginosa* has previously been established using knockout mutants. For example, deletion of the O-antigen ligase, *waaL*, responsible for ligating O-antigen molecules to the lipid A core, results in reduced twitching motility (73). How O-antigen deficiency affects twitching motility is not understood. Here, we extend these results and show that expression of different O-antigen serotypes leads to differences in twitching motility. We propose that differences in outer membrane stability as a function of different sugar compositions of the O-antigen repeat structure affect the function of T4P.

Finally, we characterized the virulence of serotype switched strains (both isogenic and clinical strains). We find that (compared to wild type PAO1), serotype O4 is associated with reduced lung colonization, and increased mouse survival during a murine acute pneumonia infection. On the other hand, serotype O12 is associated with more virulence and a higher lung colonization that is similar to wild type PAO1. We similarly characterized the survival of mice infected with clinical ST111 strains expressing serotype O4 or O12, which shows that the serotype O12 expressing strain is more virulent at similar doses of infection. We note that the clinical ST111 strains (ST111 2875 and ST111 2879) are not isogenic, and we cannot rule out contributions from genetic variations outside the O-antigen structure on the observed virulence phenotypes. Nevertheless, the clear link between serotype O12 and increased virulence in isogenic PAO1 strains, compared to serotype O4, suggests that the serotype configuration is also contributing to the virulence of ST111.

A possible explanation for this difference in virulence between serotypes is that they confer different levels of resistance to host defenses such as complement-mediated killing. We tested the serum susceptibility of the engineered *P. aeruginosa* strains and found that serotype O12 is associated with increased survival in serum, but only in the ST111 genetic background. However, we also see that the engineered ST111 strains show a higher fitness in serum compared to the wild type strain ST111 2875. As the OSA of serotype O12 contains a terminal sialic acid residue, we speculate that this could modulate host immune response or mediate specific interactions that enhance virulence (90–93). However, this requires future analysis of the host immune response to an infection with serotype switched strains.

Overall, our study reveals that *P. aeruginosa* pathogenicity traits such as adhesion, twitching motility, virulence, and tolerance to host defence molecules (i.e lysozyme) and complement-mediated killing are affected by the different sugar composition found in O4 and O12 O-antigens. We tracked these effects to differences in the biophysical properties of O4 versus O12 O-antigens, and show that expression of O4 affects the stability of the outer membrane, EDTA sensitivity, and strength of the Type IV pili. Importantly, these results provide clarifications on the potential selective advantages of the serotype switch from O4 to O12 that underlie the emergence and global dissemination of ST111 O12.

Interestingly, the serotype switch from O4 to O12 appear to be specifically associated to ST111 although serotype switches to O12 from other serotypes have been observed at low frequency in other clone types, such as ST253 and ST244 (9). It is possible that unknown interactions between host genotype and different OSA biosynthesis gene clusters restricts dissemination of the O12 OSA gene cluster across a broader range of clone types. Our observation that the OSA length regulation is dependent on the genetic background of strains used for heterologous expression (Figure 2), suggests that such interactions indeed exist. Thus, it is likely that interactions between host-specific genetic factors and OSA may affect fitness in particular environments and thereby contribute to restrict serotype switching. In support of this conclusion, recent studies have found that different, natural *Pseudomonas* isolates with identical O-antigen biosynthesis gene clusters exhibit varying sensitivities towards tailocins (94). Future investigation of interactions between host genotype and O12 OSA gene may benefit from the recombineering method described in the present work to capture, integrate and express the O12 gene cluster across different *P. aeruginosa* hosts. More broadly, the same approach may be used to explore how the 20 different O-antigen structures found in *P. aeruginosa* impacts its physiology and behaviors. Our demonstration that differences between the O4 and O12 O-antigen structures can result in substantial differences in *P. aeruginosa* phenotypes suggest that a systematic analysis of O-antigen structure/function relationship may provide new insight into the biology of *P. aeruginosa*, and contribute to refine development of anti-pseudomonas antimicrobials that targets or involves LPS O-antigen molecules.

## Author statements

### Author contributions

Conceptualization: MA & LJ. Data curation: MA. Formal Analysis: MA, DM & TOP. Funding acquisition: LJ, JBG, YFD & AB. Investigation: MA, DM, TOP, MAL & JG. Methodology: MA. Project Administration: LJ. Resources: LJ, JBG, YFD & AB. Software: MA. Supervision: LJ, JBG, YFD, LS, KZ. Validation: MA, DM, & TOP. Visualization: MA. Writing – original draft: MA. Writing – review and editing: MA, TOP, DM, JBG, KZ, and LJ.

## Acknowledgements

We acknowledge Francis A Stewart & Hailong Wang for RecET cloning strain GBdir-pir116, Shengda Zhang for RecET recombineering advice, and Janus Haagensen for the PA14 strain.

## References

1. Oliver A, Mulet X, López-Causapé C, Juan C. 2015. The increasing threat of Pseudomonas aeruginosa high-risk clones. Drug Resistance Updates 21–22:41–59.

2. del Barrio-Tofiño E, López-Causapé C, Oliver A. 2020. Pseudomonas aeruginosa epidemic high-risk clones and their association with horizontally-acquired β-lactamases: 2020 update. International Journal of Antimicrobial Agents 56:106196.

3. Treepong P, Kos VN, Guyeux C, Blanc DS, Bertrand X, Valot B, Hocquet D. 2018. Global emergence of the widespread Pseudomonas aeruginosa ST235 clone. Clinical Microbiology and Infection 24:258–266.

4. Thrane SW, Taylor VL, Freschi L, Kukavica-Ibrulj I, Boyle B, Laroche J, Pirnay J-P, Lévesque RC, Lam JS, Jelsbak L. 2015. The Widespread Multidrug-Resistant Serotype O12 Pseudomonas aeruginosa Clone Emerged through Concomitant Horizontal Transfer of Serotype Antigen and Antibiotic Resistance Gene Clusters. mBio 6:e01396–15.

5. Turton JF, Wright L, Underwood A, Witney AA, Chan YT, Al-Shahib A, Arnold C, Doumith M, Patel B, Planche TD, Green J, Holliman R, Woodford N. 2015. High-resolution analysis by whole-genome sequencing of an international lineage (Sequence Type 111) of pseudomonas aeruginosa associated with metallo-carbapenemases in the United Kingdom. Journal of Clinical Microbiology 53:2622–2631.

6. Wright LL, Turton JF, Livermore DM, Hopkins KL, Woodford N. 2015. Dominance of international ‘high-risk clones’ among metallo-β-lactamase-producing Pseudomonas aeruginosa in the UK. Journal of Antimicrobial Chemotherapy 70:103–110.

7. Pirzadian J, Persoon MC, Severin JA, Klaassen CHW, de Greeff SC, Mennen MG, Schoffelen AF, Wielders CCH, Witteveen S, van Santen-Verheuvel M, Schouls LM, Vos MC, Bode L, Troelstra A, Notermans DW, Maijer-Reuwer A, Leversteijn-van Hall MA, Kluytmans JAJW, Spijkerman IJB, van Dijk K, Halaby T, Zwart B, Diederen BMW, Voss A, Dorigo-Zetsma JW, Ott A, Oudbier JH, van der Vusse M, Vlek ALM, Buiting AGM, Paltansing S, de Man P, van Griethuysen AJ, den Reijer M, van Trijp M, van Elzakker EPM, Muller AE, van der Linden MPM, van Rijn M, Wolfhagen MJHM, Waar K, Schneeberger P, Silvis W, Schulin T, Damen M, Dinant S, van Mens SP, Melles DC, Cohen Stuart JWT, van Ogtrop ML, Overdevest ITMA, van Dam A, Wertheim H, Frénay HME, Sinnige JC, Mattsson EE, Bosboom RW, Stam A, de Jong E, Roescher N, Heikens E, Steingrover R, Bathoorn E, Trienekens TAM, van Dam DW, de Brauwer EIGB, Stals FS. 2021. National surveillance pilot study unveils a multicenter, clonal outbreak of VIM-2-producing Pseudomonas aeruginosa ST111 in the Netherlands between 2015 and 2017. Scientific Reports 2021 11:1 11:1–10.

8. Guzvinec M, Izdebski R, Butic I, Jelic M, Abram M, Koscak I, Baraniak A, Hryniewicz W, Gniadkowski M, Andrasevic AT. 2014. Sequence types 235, 111, and 132 Predominate among multidrug-resistant Pseudomonas aeruginosa clinical isolates in croatia. Antimicrobial Agents and Chemotherapy 58:6277–6283.

9. Anbo M, Jelsbak L. 2023. A bittersweet fate: detection of serotype switching in Pseudomonas aeruginosa. Microbial Genomics 9:000919.

10. Pitt TL, Livermore DM, Pitcher D, Vatopoulos AC, Legakis NJ. 1989. Multiresistant serotype O 12 Pseudomonas aeruginosa: evidence for a common strain in Europe. Epidemiology & Infection 103:565–576.

11. Huszczynski SM, Lam JS, Khursigara CM. 2020. The Role of Pseudomonas aeruginosa Lipopolysaccharide in Bacterial Pathogenesis and Physiology. Pathogens 9:6.

12. Lam J, Taylor V, Islam S, Hao Y, Kocíncová D. 2011. Genetic and Functional Diversity of Pseudomonas aeruginosa Lipopolysaccharide. Frontiers in Microbiology 2.

13. Peterson AA, Hancock REW, McGroarty EJ. 1985. Binding of polycationic antibiotics and polyamines to lipopolysaccharides of Pseudomonas aeruginosa. Journal of Bacteriology 164:1256– 1261.

14. Hancock RE. 1984. Alterations in outer membrane permeability. Annual review of microbiology 38:237–264.

15. Fernández L, Álvarez-Ortega C, Wiegand I, Olivares J, Kocíncová D, Lam JS, Martínez JL, Hancock REW. 2013. Characterization of the polymyxin B resistome of Pseudomonas aeruginosa. Antimicrobial Agents and Chemotherapy 57:110–119.

16. Alvarez-Ortega C, Wiegand I, Olivares J, Hancock REW, Martínez JL. 2010. Genetic determinants involved in the susceptibility of Pseudomonas aeruginosa to β-lactam antibiotics. Antimicrobial Agents and Chemotherapy 54:4159–4167.

17. Knirel YA, Bystrova OV, Kocharova NA, Zähringer U, Pier GB. 2006. Conserved and variable structural features in the lipopolysaccharide of Pseudomonas aeruginosa. Journal of endotoxin research 12:324–336.

18. LIU PV, MATSUMOTO H, KUSAMA H, BERGAN T. 1983. Survey of Heat-Stable, Major Somatic Antigens of Pseudomonas aeruginosa†. International Journal of Systematic and Evolutionary Microbiology 33:256–264.

19. Liu PV, Wang S. 1990. Three new major somatic antigens of Pseudomonas aeruginosa. Journal of Clinical Microbiology 28:922–925.

20. Knirel YA, Vinogradov EV, Kocharova NA, Paramonov NA, Kochetkov NK, Dmitriev BA, Stanislavsky ES, Lanyi B. 1988. The structure of O-specific polysaccharides and serological classification of Pseudomonas aeruginosa. Acta Microbiologica Hungarica 35:3–24.

21. Cryz SJ, Pitt TL, Furer E, Germanier R. 1984. Role of lipopolysaccharide in virulence of Pseudomonas aeruginosa. Infection and Immunity 44:508–513.

22. Dötsch A, Becker T, Pommerenke C, Magnowska Z, Jänsch L, Häussler S. 2009. Genomewide identification of genetic determinants of antimicrobial drug resistance in Pseudomonas aeruginosa. Antimicrobial Agents and Chemotherapy 53:2522–2531.

23. Engels W, Endert J, Kamps MAF, Van Boven CPA. 1985. Role of lipopolysaccharide in opsonization and phagocytosis of Pseudomonas aeruginosa. Infection and Immunity 49:182–189.

24. Pitt TL. 1988. Epidemiological typing ofPseudomonas aeruginosa. Eur J Clin Microbiol Infect Dis 7:238–247.

25. Herrero M, De Lorenzo V, Timmis KN. 1990. Transposon vectors containing non-antibiotic resistance selection markers for cloning and stable chromosomal insertion of foreign genes in gram-negative bacteria. Journal of Bacteriology 172:6557–6567.

26. Wang H, Li Z, Jia R, Hou Y, Yin J, Bian X, Li A, Müller R, Stewart AF, Fu J, Zhang Y. 2016. RecET direct cloning and Redαβ recombineering of biosynthetic gene clusters, large operons or single genes for heterologous expression. Nature Protocols 2016 11:7 11:1175–1190.

27. Anbo M, Chichkova MAT, Gençay YE, Salazar A, Jeannot K, Jelsbak L. 2023. Whole-Genome Sequencing of 11 High-Risk Clone ST111 Pseudomonas aeruginosa Isolates from French Hospitals. Microbiology Resource Announcements 12:e00091–23.

28. Dean CR, Goldberg JB. 2002. Pseudomonas aeruginosa galU is required for a complete lipopolysaccharide core and repairs a secondary mutation in a PA103 (serogroup O11) wbpM mutant. FEMS Microbiology Letters 210:277–283.

29. Rau MH, Hansen SK, Johansen HK, Thomsen LE, Workman CT, Nielsen KF, Jelsbak L, Høiby N, Yang L, Molin S. 2010. Early adaptive developments of Pseudomonas aeruginosa after the transition from life in the environment to persistent colonization in the airways of human cystic fibrosis hosts. Environmental Microbiology 12:1643–1658.

30. Kessler B, de Lorenzo V, Timmis KN. 1992. A general system to integratelacZ fusions into the chromosomes of gram-negative eubacteria: regulation of thePm promoter of theTOL plasmid studied with all controlling elements in monocopy. Molecular and General Genetics MGG 1992 233:1 233:293–301.

31. Choi KH, Gaynor JB, White KG, Lopez C, Bosio CM, Karkhoff-Schweizer RAR, Schweizer HP. 2005. A Tn7-based broad-range bacterial cloning and expression system. Nature Methods 2005 2:6 2:443– 448.

32. Yang L, Hengzhuang W, Wu H, Damkiær S, Jochumsen N, Song Z, Givskov M, Høiby N, Molin S. 2012. Polysaccharides serve as scaffold of biofilms formed by mucoid Pseudomonas aeruginosa. FEMS Immunology and Medical Microbiology 65:366–376.

33. Lambertsen L, Sternberg C, Molin S. 2004. Mini-Tn7 transposons for site-specific tagging of bacteria with fluorescent proteins. Environmental Microbiology 6:726–732.

34. Zobel S, Benedetti I, Eisenbach L, de Lorenzo V, Wierckx N, Blank LM. 2015. Tn7-Based Device for Calibrated Heterologous Gene Expression in Pseudomonas putida. ACS Synth Biol 4:1341–1351.

35. Hoang TT, Kutchma AJ, Becher A, Schweizer HP. 2000. Integration-Proficient Plasmids for Pseudomonas aeruginosa: Site-Specific Integration and Use for Engineering of Reporter and Expression Strains. Plasmid 43:59–72.

36. Hmelo LR, Borlee BR, Almblad H, Love ME, Randall TE, Tseng BS, Lin C, Irie Y, Storek KM, Yang JJ, Siehnel RJ, Howell PL, Singh PK, Tolker-Nielsen T, Parsek MR, Schweizer HP, Harrison JJ. 2015. Precision-engineering the Pseudomonas aeruginosa genome with two-step allelic exchange. Nature Protocols 2015 10:11 10:1820–1841.

37. Chen S, Zhou Y, Chen Y, Gu J. 2018. Fastp: An ultra-fast all-in-one FASTQ preprocessor, p. i884– i890. In Bioinformatics. Oxford Academic.

38. Li H. 2023. Seqtk: Toolkit for processing sequences in FASTA/Q formats. https://github.com/lh3/seqtk.

39. Wick R. rrwick/Porechop: adapter trimmer for Oxford Nanopore reads.

40. Wick R. rrwick/Filtlong: quality filtering tool for long reads.

41. Wick RR, Judd LM, Gorrie CL, Holt KE. 2017. Unicycler: Resolving bacterial genome assemblies from short and long sequencing reads. PLOS Computational Biology 13:e1005595.

42. Tatusova T, DiCuccio M, Badretdin A, Chetvernin V, Nawrocki EP, Zaslavsky L, Lomsadze A, Pruitt KD, Borodovsky M, Ostell J. 2016. NCBI prokaryotic genome annotation pipeline. Nucleic Acids Research 44:6614–6624.

43. O’Toole GA, Pratt LA, Watnick PI, Newman DK, Weaver VB, Kolter R. 1999. [6] Genetic approaches to study of biofilms. Methods in Enzymology 310:91–109.

44. Petrauskas AA, Kolovanov EA. 2000. ACD/Log P method description. Perspectives in Drug Discovery and Design 19:99–116.

45. Ayres HM, Furr JR, Russell AD. 1998. Effect of divalent cations on permeabilizer-induced lysozyme lysis of Pseudomonas aeruginosa. Letters in Applied Microbiology 27:372–374.

46. Turnbull L, Whitchurch CB. 2014. Motility assay: twitching motility. Methods in Molecular Biology (Clifton, NJ) 1149:73–86.

47. Beaussart A, Baker AE, Kuchma SL, El-Kirat-Chatel S, Otoole GA, Dufrêne YF. 2014. Nanoscale adhesion forces of Pseudomonas aeruginosa type IV pili. ACS Nano 8:10723–10733.

48. Beaussart A, El-Kirat-Chatel S, Herman P, Alsteens D, Mahillon J, Hols P, Dufrêne YF. 2013. Single-Cell Force Spectroscopy of Probiotic Bacteria. Biophysical Journal 104:1886–1892.

49. Viljoen A, Vercellone A, Chimen M, Gaibelet G, Mazères S, Nigou J, Dufrêne YF. 2023. Nanoscale clustering of mycobacterial ligands and DC-SIGN host receptors are key determinants for pathogen recognition. Science Advances 9:eadf9498.

50. Webster SS, Mathelié-Guinlet M, Verissimo AF, Schultz D, Viljoen A, Lee CK, Schmidt WC, Wong GCL, Dufrêne YF, O’Toole GA. 2022. Force-Induced Changes of PilY1 Drive Surface Sensing by Pseudomonas aeruginosa. mBio 13:e03754–21.

51. Moustafa DA, Scarff JM, Garcia PP, Cassidy SKB, DiGiandomenico A, Waag DM, Inzana TJ, Goldberg JB. 2015. Recombinant Salmonella Expressing Burkholderia mallei LPS O Antigen Provides Protection in a Murine Model of Melioidosis and Glanders. PLoS One 10:e0132032.

52. Kassambara A. 2020. “ggplot2” Based Publication Ready Plots [R package ggpubr version 0.4.0].

53. Therneau TM, until 2009) maintainer TL (original S-\textgreaterR port, R., Elizabeth A, Cynthia C. 2023. survival: Survival Analysis.

54. Valero-Mora PM. 2010. ggplot2: Elegant Graphics for Data Analysis. Springer-Verlag New York. https://ggplot2.tidyverse.org.

55. Hadley W, Jim H, Jennifer B. 2021. Read Rectangular Text Data [R package readr version 2.1.1].

56. Garnier, Simon, Ross, Noam, Rudis, Robert, Camargo, Pedro A, Sciaini, Marco, Scherer, Cédric. 2021. {viridis} - Colorblind-Friendly Color Maps for R.

57. Horikoshi M, Tang [aut Y, cre, Dickey A, Grenié M, Thompson R, Selzer L, Strbenac D, Voronin K, Pulatov D. 2023. ggfortify: Data Visualization Tools for Statistical Analysis Results.

58. Kintz E, Scarff JM, DiGiandomenico A, Goldberg JB. 2008. Lipopolysaccharide O-Antigen Chain Length Regulation in Pseudomonas aeruginosa Serogroup O11 Strain PA103. Journal of Bacteriology 190:2709–2716.

59. Makin SA, Beveridge TJ. 1996. The influence of A-band and B-band lipopolysaccharide on the surface characteristics and adhesion of Pseudomonas aeruginosa to surfaces. Microbiology 142:299–307.

60. Hermansson M. 1999. The DLVO theory in microbial adhesion. Colloids and Surfaces B: Biointerfaces 14:105–119.

61. Bos R, Van Der Mei HC, Busscher HJ. 1999. Physico-chemistry of initial microbial adhesive interactions - Its mechanisms and methods for study. FEMS Microbiology Reviews 23:179–230.

62. Poortinga AT, Bos R, Norde W, Busscher HJ. 2002. Electric double layer interactions in bacterial adhesion to surfaces. Surface Science Reports 47:1–32.

63. van Loosdrecht MCM, Lyklema J, Norde W, Zehnder AJB. 1989. Bacterial adhesion: A physicochemical approach. Microbial Ecology 17:1–15.

64. Lerman MJ, Lembong J, Muramoto S, Gillen G, Fisher JP. 2018. The Evolution of Polystyrene as a Cell Culture Material. Tissue Engineering - Part B: Reviews 24:359–372.

65. Cox Jr. ST, Eagon RG. 1968. Action of ethylenediaminetetraacetic acid, tris(hydroxymethyl)aminomethane, and lysozyme on cell walls of Pseudomonas aeruginosa. Canadian Journal of Microbiology 14:913–922.

66. Clifton LA, Skoda MWA, Le Brun AP, Ciesielski F, Kuzmenko I, Holt SA, Lakey JH. 2015. Effect of Divalent Cation Removal on the Structure of Gram-Negative Bacterial Outer Membrane Models. Langmuir 31:404–412.

67. Powell KJ, Brown PL, Byrne RH, Gajda T, Hefter G, Sjöberg S, Wanner H. 2005. Chemical speciation of environmentally significant heavy metals with inorganic ligands part 1: The Hg2+-Cl-, OH-, CO32-, so42-, and PO 43-aqueous systems (IUPAC technical report). Pure and Applied Chemistry 77:739–800.

68. King JD, Mulrooney EF, Vinogradov E, Kneidinger B, Mead K, Lam JS. 2008. lfnA from Pseudomonas aeruginosa O12 and wbuX from Escherichia coli O145 encode membrane-associated proteins and are required for expression of 2,6-dideoxy-2-acetamidino-L-galactose in lipopolysaccharide O antigen. Journal of Bacteriology 190:1671–1679.

69. Knirel YA, Vinogradov EV, Shashkov AS, Dmitriev BA, Kochetkov NK, Stanislavsky ES, Mashilova GM. 1987. Somatic antigens of Pseudomonas aeruginosa The structure of the O-specific polysaccharide chain of the lipopolysaccharide from P aeruginosa O13 (Lányi). European Journal of Biochemistry 163:627–637.

70. Dufrêne YF, Viljoen A, Mignolet J, Mathelié-Guinlet M. 2021. AFM in cellular and molecular microbiology. Cellular Microbiology 23:e13324.

71. Burrows LL. 2012. Pseudomonas aeruginosa twitching motility: Type IV pili in action. Annual Review of Microbiology 66:493–520.

72. Mattick JS. 2002. Type IV Pili and Twitching Motility. Annual Review of Microbiology 56:289–314.

73. Abeyrathne PD, Daniels C, Poon KKH, Matewish MJ, Lam JS. 2005. Functional characterization of WaaL, a ligase associated with linking O-antigen polysaccharide to the core of Pseudomonas aeruginosa lipopolysaccharide. Journal of Bacteriology 187:3002–3012.

74. Ivanov IE, Kintz EN, Porter LA, Goldberg JB, Burnham NA, Camesano TA. 2011. Relating the physical properties of pseudomonas aeruginosa lipopolysaccharides to virulence by atomic force microscopy. Journal of Bacteriology 193:1259–1266.

75. Lau PCY, Lindhout T, Beveridge TJ, Dutcher JR, Lam JS. 2009. Differential lipopolysaccharide core capping leads to quantitative and correlated modifications of mechanical and structural properties in Pseudomonas aeruginosa biofilms. Journal of Bacteriology 191:6618–6631.

76. Abu-Lail LI, Liu Y, Atabek A, Camesano TA. 2007. Quantifying the adhesion and interaction forces between Pseudomonas aeruginosa and natural organic matter. Environmental Science and Technology 41:8031–8037.

77. Atabek A, Camesano TA. 2007. Atomic force microscopy study of the effect of lipopolysaccharides and extracellular polymers on adhesion of Pseudomonas aeruginosa. Journal of bacteriology 189:8503–8509.

78. Williams V, Fletcher M. 1996. Pseudomonas fluorescens adhesion and transport through porous media are affected by lipopolysaccharide composition. Applied and Environmental Microbiology 62:100–104.

79. Pringle JH, Fletcher M. 1983. Influence of Substratum Wettability on Attachment of Freshwater Bacteria to Solid Surfaces. Applied and Environmental Microbiology 45:811–817.

80. Burrows LL, Lam JS. 1999. Effect of wzx (rfbX) mutations on A-band and B-band lipopolysaccharide biosynthesis in Pseudomonas aeruginosa O5. Journal of Bacteriology 181:973–980.

81. Rocchetta HL, Burrows LL, Pacan JC, Lam JS. 1998. Three rhamnosyltransferases responsible for assembly of the A-band D-rhamnan polysaccharide in Pseudomonas aeruginosa: A fourth transferase, WbpL, is required for the initiation of both A-band and B-band lipopolysaccharide synthesis. Molecular Microbiology 28:1103–1119.

82. Burrows LL, Urbanic RV, Lam JS. 2000. Functional conservation of the polysaccharide biosynthetic protein WbpM and its homologues in Pseudomonas aeruginosa and other medically significant bacteria. Infection and immunity 68:931–936.

83. Bélanger M, Burrows LL, Lam JS. 1999. Functional analysis of genes responsible for the synthesis of the B-band O antigen of pseudomonas aeruginosa serotype O6 lipopolysaccharide. Microbiology 145:3505–3521.

84. Dean CR, Franklund CV, Retief JD, Coyne MJ, Hatano K, Evans DJ, Pier GB, Goldberg JB. 1999. Characterization of the serogroup O11 O-antigen locus of Pseudomonas aeruginosa PA103. Journal of Bacteriology 181:4275–4284.

85. Hao Y, Murphy K, Lo RY, Khursigara CM, Lam JS. 2015. Single-nucleotide polymorphisms found in the migA and wbpX glycosyltransferase genes account for the intrinsic lipopolysaccharide defects exhibited by Pseudomonas aeruginosa PA14. Journal of Bacteriology 197:2780–2791.

86. Azimi S, Thomas J, Cleland SE, Curtis JE, Goldberg JB, Diggle SP. 2021. O-Specific Antigen-Dependent Surface Hydrophobicity Mediates Aggregate Assembly Type in Pseudomonas aeruginosa. mBio 12:10.1128/mbio.00860-21.

87. Ulrich N, Goss KU, Ebert A. 2021. Exploring the octanol–water partition coefficient dataset using deep learning techniques and data augmentation. Communications Chemistry 2021 4:1 4:1–10.

88. Vimr ER, Kalivoda KA, Deszo EL, Steenbergen SM. 2004. Diversity of Microbial Sialic Acid Metabolism. Microbiology and Molecular Biology Reviews 68:132.

89. Langley S, Beveridge TJ. 1999. Effect of O-side-chain-lipopolysaccharide chemistry on metal binding. Applied and Environmental Microbiology 65:489–498.

90. Severi E, Hood DW, Thomas GH. 2007. Sialic acid utilization by bacterial pathogens. Microbiology 153:2817–2822.

91. Chang YC, Olson J, Beasley FC, Tung C, Zhang J, Crocker PR, Varki A, Nizet V. 2014. Group B Streptococcus Engages an Inhibitory Siglec through Sialic Acid Mimicry to Blunt Innate Immune and Inflammatory Responses In Vivo. PLoS Pathogens 10:e1003846.

92. Rasko DA, Keelan M, Wilson TJM, Taylor DE. 2001. Lewis Antigen Expression by Helicobacter pylori. The Journal of Infectious Diseases 184:315–321.

93. Varki A, Gagneux P. 2012. Multifarious roles of sialic acids in immunity. Annals of the New York Academy of Sciences 1253:16–36.

94. Carim S, Azadeh AL, Kazakov AE, Price MN, Walian PJ, Lui LM, Nielsen TN, Chakraborty R, Deutschbauer AM, Mutalik VK, Arkin AP. 2021. Systematic discovery of pseudomonad genetic factors involved in sensitivity to tailocins. 8. ISME J 15:2289–2305.

